# Bioengineered embryoids mimic post-implantation development *in vitro*

**DOI:** 10.1101/2021.01.10.426096

**Authors:** Mehmet U. Girgin, Nicolas Broguiere, Sylke Hoehnel, Nathalie Brandenberg, Bastien Mercier, Alfonso Martinez Arias, Matthias P. Lutolf

## Abstract

The difficulty of studying post-implantation development in mammals has sparked a flurry of activity to develop *in vitro* models, termed embryoids, based on self-organizing pluripotent stem cells. Previous approaches to derive embryoids either lack the physiological morphology and signaling interactions, or are not yet optimal for modeling post-gastrulation development. Here, we report a bioengineering-inspired approach aimed at addressing this gap. A high-throughput cell aggregation approach was employed to simultaneously coax mouse embryonic stem cells (ESCs) into hundreds of uniform epiblast-like (EPI) aggregates in a solid matrix-free manner. When co-cultured with mouse trophoblast stem cell (TSC) aggregates, the resulting hybrid structures initiate gastrulation-like events and undergo axial morphogenesis to yield structures, termed *EpiTS embryoids*, with a pronounced anterior development, including brain-like regions. We identify the presence of an epithelium in EPI aggregates as the major determinant for the axial morphogenesis and anterior development seen in *EpiTS embryoids*. Our results demonstrate the potential of *EpiTS embryoids* to study peri-gastrulation development *in vitro*.

## Introduction

Recapitulating early mammalian development *in vitro* is an important challenge that could overcome the experimental bottleneck created by intrauterine development and assist in the reduction of the use of animals in research^1^. The control of the differentiation potential of ESCs has been a central feature in these efforts. Constraining the differentiating cells within micropatterned substrates controls the stochastic differentiation of cells in adherent culture and results in patterns that resemble the organization of germ layers in the embryo^2,3^. Three-dimensional embryoid models trigger selforganization programs that mimic much of the early development of the embryo and overcome the heterogeneous and heterochronic patterns characteristic of Embryoid Bodies (EBs)^4^. In particular, gastruloids^5^, aggregates of defined numbers of ESCs, develop derivatives of all germ layers with spatiotemporal patterns characteristic of embryos even though they lack clear brain structures^6^. Gastruloids have proven useful tools to explore the consequences of gastrulation in the absence of extraembryonic tissues, such as formation of cardiac primordia^7^ or somite-like structures^8,9^. Other embryoid models have explored interactions between embryonic and extraembryonic tissues during early development by taking advantage of the existence of trophoblast (TSC) and extraembryonic endoderm (XEN) stem cells to recreate the early conceptus. Co-culture of TSCs and XEN cells with mouse ESCs in Matrigel results in a suite of structures, including blastoids^10^, ETS^11^- and ETX^12^-embryos, that recapitulate events and interactions of the pre-gastrulation embryo. A challenge with ETS- and ETX-embryos has been the reliance on the self-organizing activity of the cellular compartments that results in their stochastic occurrence; only ~ 20% of the starting aggregates will form patterned structures^11^, which limits their broader applicability. Furthermore, their developmental potential is not yet clear, as there are no reports of their development beyond early gastrulation stages^11^. Altogether, these pioneering *in vitro* models of early embryo development highlight the remarkable capacity of embryonic and extraembryonic cells to organize themselves but thus far have not yet been ideal to explore the fate and derivatives of cells in the emerging structures.

Here, we used a bioengineering approach combining features of ETS embryos and gastruloids to build an embryoid model that could more realistically capture the events leading to and resulting from gastrulation. We hypothesized that to achieve this, it would be essential to optimize the starting culture conditions in multicellular aggregates to assemble, in the same embryoid, an anterior neuroepithelium with a posterior *T/Bra* expressing cell ensemble. We used arrays of cell-repellent hydrogel microwells to separately generate epithelialized or non-epithelialized epiblast-like (EPI) aggregates, as well as TSC aggregates mimicking extraembryonic ectoderm, in a scalable manner. When assembled together in low-attachment wells in serum-free medium, EPI and TSC aggregates rapidly merged and underwent symmetry breaking similar to gastrulating embryos, as demonstrated by polarized and restricted *Brachyury* (*T/Bra*) expression. Importantly, in these structures, *T/Bra* expression dynamics was strictly dependent on the epithelial architecture of the EPI aggregates. Subsequently, hybrid EPI/TSC structures, termed *EpiTS embryoids*, underwent axial morphogenesis to display patterning along anterior-posterior, dorsal-ventral and medio-lateral axes. Strikingly, we observed that *embryoids* formed from non-epithelialized EPI aggregates predominantly generated mesendodermal tissue, whereas epithelialized ones formed cell types that are present in the developing midbrain/hindbrain. Our approach enables the generation of various tissues in a stereotyped and scalable manner, with independent modulation of physical (e.g., size, epithelial architecture) and biological (e.g., provision of signaling molecules) properties of EPI and TSC aggregates, such as to systematically parse out the role of these parameters in promoting key steps in embryogenesis.

## Results

### Scalable formation of homogeneous EPI aggregates

We aggregated ESCs in round-bottom microwell arrays composed of non-adhesive PEG hydrogels^13^ in epiblast induction medium comprising Activin-A, bFGF and KSR, supplemented with low percentage (3%) Matrigel to induce epithelialization^14^ (**Fig. 1a**). The microwells allowed us to titrate the average number of cells seeded in each well, resulting in aggregates of defined size (**Fig. 1b, Supplementary Fig. 1a,** upper panel). The starting number of cells determined the size of the aggregate at 72 h of culture: an average of ~25 ESCs per microwell yielded aggregates with ~180 μm diameter, whereas seeding ~100 cells reached a diameter of ~230 μm (**Fig. 1c**). These aggregates featured a single lumen surrounded by an *E-cadherin*-positive polarized epithelium, displaying apical *Pdx* and *Par6* expression (**Fig. 1d-f,** lower panel; **Supplementary Fig. 1c**). Smaller aggregates exhibited a discontinuous apical expression of *aPKC* surrounded by a multi-layered *E-cadherin-positive* epithelium and multiple *F-actin* labeled cavities, suggesting poor epithelialization and incomplete lumenization (**Fig. 1d-f,** upper panel). These results demonstrate that a critical aggregate size needs to be reached to form an apico-basally polarized epithelium with a central lumen, a phenomenon reminiscent of epiblast maturation^15^. When cells were aggregated in the absence of Matrigel, we detected increased cell shedding (**Supplementary Fig. 1a,** lower panel, white arrows) but no significant difference in aggregate diameter (**Supplementary Fig. 1b**). However, these aggregates composed of *E-cadherin*-positive cells demonstrated inverted *Factin* polarity and featured multiple *Pdx*-positive and *Par6*-positive foci with no clear epithelium or lumen (**Fig. 1g, Supplementary Fig. 1d**). These results show that polarized epithelial aggregates of defined size can be readily generated from ESCs using hydrogel microwell arrays, and confirmed that the provision of Matrigel is critical for their lumenization and epithelialization^16^.

**Figure 1:**
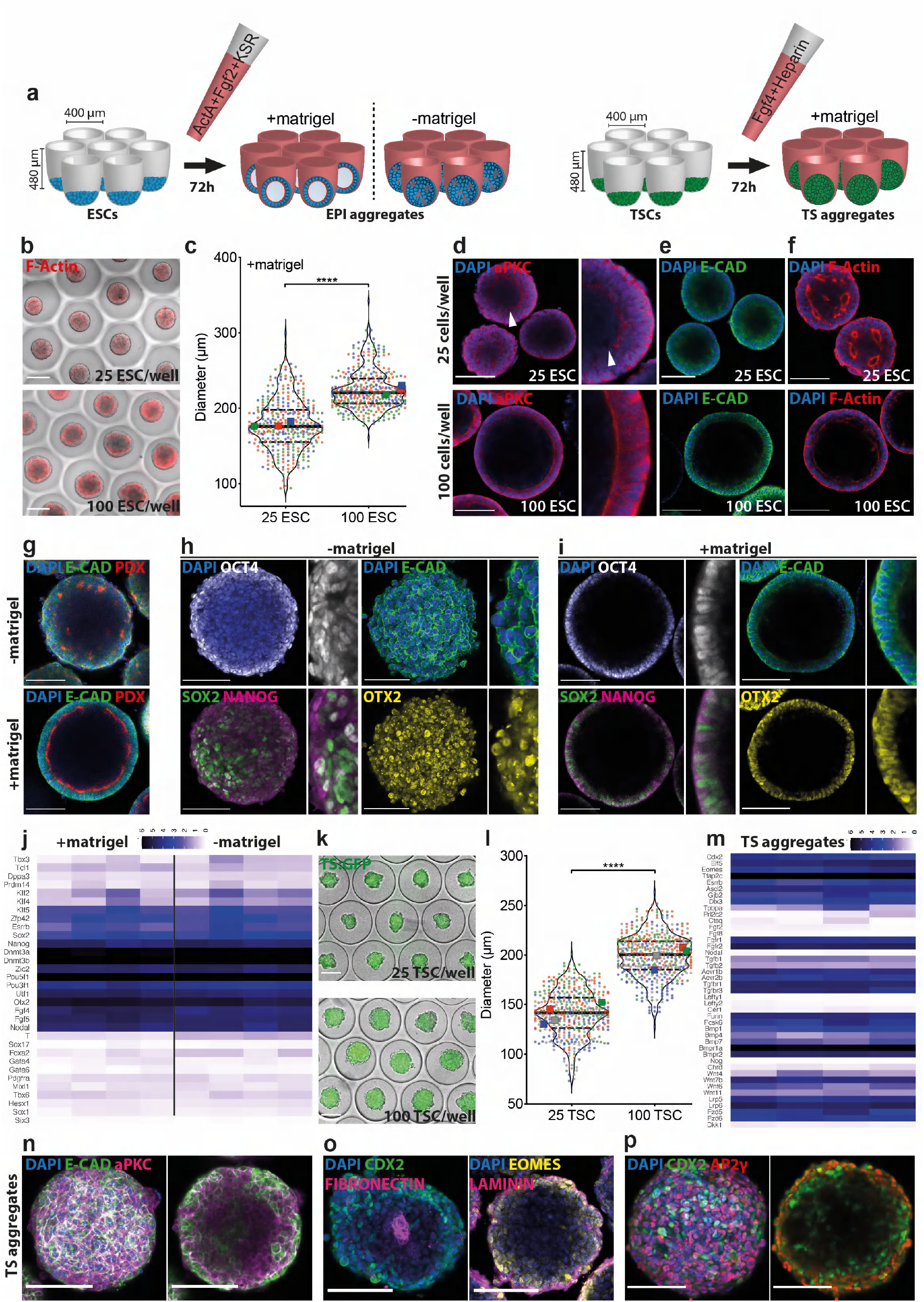
Formation and characterization of EPI and TS aggregates. **a)** Schematic representation of the workflow showing aggregation of ESCs and TSCs on PEG microwells. **b)** Representative images showing EPI aggregates on microwells at 72 h formed from 25 cells/well (top) or 100 cells/well (bottom) showing F-actin staining by phalloidin. Scale bars: 200μm. **c)** Comparing minimum ferret diameters of EPI aggregates at 72 h formed from 25 or 100 cells/well. For 25 ESC and 100 ESC conditions, total number of embryoids analyzed were 362 and 361, respectively. Data is collected from three biologically independent experiments. Large symbols indicate mean values of each replicate. Black lines indicate median and quartiles. **d-f)** Confocal images of EPI aggregates formed from 25 cells/well or 100 cells/well fixed at 72 h and stained for *aPKC* **(d)***, E-cadherin* **(e)** and *F-actin* by phalloidin **(f)**. Nuclei were stained with DAPI. Note multi-layered epithelium in EPI aggregates formed from 25 cells/well showing discontinuous *aPKC* staining (white arrows) compared to single-layer epithelium in EPI aggregates formed from 100 cells/well showing continuous *aPKC* signal. Scale bars: 100μm. **g)** Confocal images of aggregates formed from 100 cells/well with or without Matrigel fixed at 72 h and stained for *E-cadherin* and *Podocalxyin*. Nuclei were stained with DAPI. Scale bars: 100μm. **h,i)** Confocal images of EPI aggregates formed from 100 cells/well without (**h)** or with (**i)** Matrigel, fixed at 72 h showing expression of *Otx2, Oct4, Sox2, Nanog* and *E-cadherin*. Nuclei were stained with DAPI. Scale bars: 100μm. **j)** Bulk RNA sequencing analysis of epithelialized and non-epithelialized EPI aggregates at 72 h formed from 100 cells/well, showing expression levels of pluripotency, epiblast-specific and early differentiation genes. Data is collected from four biologically independent experiments. **k)** Representative images showing TS aggregates on microwells at 72 h formed from 25 cells/well (top) or 100 cells/well (bottom) showing ubiquitous GFP expression. Scale bars: 200μm. **l)** Comparing minimum ferret diameters of TS aggregates at 72 h formed from 25 or 100 cells/well. For 25 TSC and 100 TSC conditions, total number of embryoids analyzed were 455 and 478, respectively. Data is collected from four biologically independent experiments. Large symbols indicate mean values of each replicate. Black lines indicate median and quartiles. **m)** Bulk RNA sequencing analysis of TS aggregates at 72 h formed from 100 cells/well, showing expression levels of stem cell and differentiation markers as well as key pathway agonists/antagonists. Data is collected from four biologically independent experiments. **n-p)** Confocal images of TS aggregates formed from 100 cells/well, fixed at 72 h showing expression of *E-cadherin* and *aPKC* **(n)**, *Cdx2, Fibronectin, Laminin* and *Eomes* **(o)**, *Cdx2* and *Ap2γ* **(p)**. Nuclei were stained with DAPI. Scale bars: 100μm. For statistical analysis, two-tailed unpaired Student’s t-test **(c,l)** or multiple t-tests followed by Holm-Sidak multiple comparison test **(j,m)** were performed. Following P-value style was used: P****<0.0001, P***<0.0002, P**<0.0021, P*<0.0332.

Of note, affecting cytoskeletal activity by inhibiting stress fiber formation (Y-27632) and disrupting the actin network (CK-666) resulted in loss of a continuous apicobasally polarized epithelium. Interestingly, inducing stress fiber formation with LPA did not have any effect on epithelialization (**Supplementary Fig. 2a,b**). Moreover, blocking actin-myosin interaction (Blebbistatin), myosin activity (ML-7 and Calcyculin-A) or actin polarization dynamics (Cytochalasin-D/Latrunculin-A and Jasplakinolide) prevented formation of compact aggregates and, in some cases (ML7 and Cyto-D), epithelialization. Interestingly, treatment with the *PTEN* inhibitor BpV showed a similar effect, in line with previous reports highlighting the role of *PTEN* in epiblast polarization^17^. These results suggested that actin polymerization and actin-myosin interaction play crucial roles in the polarization and epithelialization of EPI aggregates.

To test the identity of the aggregates after 72 h, we performed immunostaining for pluripotency and epiblast markers. In the absence of Matrigel, the aggregates showed a uniform expression of *Otx2* and *E-cadherin* (**Fig. 1h**). Interestingly, *Sox2* expression was detected in a polarized fashion in cells that showed high WNT and low TGF-β activity (**Fig. 1h, Supplementary Fig. 3a**), suggestive of an axis establishment similar to 48-72 h *gastruloids*^18^. Epithelialized aggregates were comprised of *E-cadherin*+ cells organized in a columnar epithelium and were uniformly positive for *Oct4* and *Otx2*. Expression of *Sox2* and *Nanog* were detected at lower levels, often in a salt-and-pepper fashion. In general, *Sox2+* cells in epithelialized aggregates demonstrated high WNT and low TGF-β activity (**Fig. 1i, Supplementary Fig. 3b**). Bulk RNA sequencing revealed low expression levels for naïve pluripotency factors *Klf2/4, Dppa3, Tbx3* and higher levels of epiblast-specific genes *Fgf5, Otx2, Utf1* in EPI aggregates. At this stage, transcripts that mark further differentiated states such as *T, Sox1* or *Sox17* were not upregulated (**Fig. 1i**). Altogether, these observations suggested that ESCs could be coaxed into EPI aggregates that morphologically and transcriptionally resemble post-implantation epiblast.

### Scalable formation of homogeneous TSC aggregates

In the early embryo, the epiblast develops in conjunction with extraembryonic tissues, and signaling from the extraembryonic ectoderm is involved in specifying the onset of gastrulation^19^. To mimic these interactions, we used the same hydrogel microwell array technology to generate TSC aggregates (**Fig. 1k**). On average, TSC aggregates were slightly smaller than EPI aggregates but their size at 72 h was also found to be dependent on the initial cell seeding concentration: aggregates composed of 25 cells or 100 cells reached a diameter of ~140μm or ~200μm, respectively (**Fig. 1l**). We detected expression of *Cdx2, Elf5* and *Eomes* transcripts and high levels of *Tfap2c* (**Fig. 1m**), suggesting that aggregated TSCs maintain stem-cell identity and get primed for differentiation^20^. At this timepoint, genes expressed in more differentiated cell types such as spongiotrophoblast or giant cells were detected at relatively low levels (**Fig 1m**), demonstrating that TSC aggregates have acquired an intermediate state, likely corresponding to extraembryonic ectoderm^21^. Importantly, TSC aggregates expressed several receptors for FGF and TGF-β pathways but not many of the ligands, in line with previous reports suggesting TSCs as source of Fgf4 and Nodal for embryonic development^22^. For BMP and WNT pathways, we could detect both receptor and ligand expression in TSC aggregates^23^.

Unlike EPI aggregates, TSC aggregates grown in the presence of low percentage Matrigel did not lumenize (**Fig. 1n**). Immunostaining showed that TSC aggregates could interact with the surrounding *laminin*-based extracellular matrix (ECM) and deposited *fibronectin* at the aggregate core (**Fig. 1o**). Expression of the stem cell markers *Cdx2, Tfap2c* and *Eomes* were primarily maintained on the periphery, suggesting initiation of differentiation from the core of the TSC aggregates^24^ (**Fig. 1o,p**).

### Formation of *EpiTS embryoids*

Next, we assembled the EPI and TSC aggregates into structures that we termed *EpiTS embryoids*. Individual EPI and TSC aggregates were transferred at 72 – 75 h to U-bottom low-attachment wells of a 96-well plate, where they fused together within a few hours (**Fig. 2a,** top panel). Time-lapse imaging revealed that by 120 h, *T/Bra* expression appeared at the aggregate interface. The size of EPI and TSC aggregates strongly influenced the timing of *T/Bra* expression (**Fig. 2b**). *Embryoids* composed of smaller EPI aggregates initiated *T/Bra* expression generally before 110 h; by 120 h, all *embryoids* had *T/Bra*-expressing cells regardless of the size of TSC aggregates they were fused to (**Fig. 2a,** bottom panel). On the other hand, *EpiTS embryoids* formed from larger EPI aggregates showed a delayed onset of *T/Bra* expression (**Fig. 2b**), with 50-70% being *T/Bra*-positive at 120 h **(Fig. 2a,** bottom panel). In addition to differences in timing, we observed a strong dependence of the initial EPI aggregate size on the *T/Bra* expression domain (**Fig. 2c**). At 120 h, *embryoids* formed from smaller EPI aggregates acquired a dispersed expression of *T/Bra*, covering almost the entire EPI compartment, whereas *EpiTS embryoids* formed from bigger EPI aggregates featured a restricted *T/Bra* expression **(Supplementary Movies 1-4)**.

**Figure 2:**
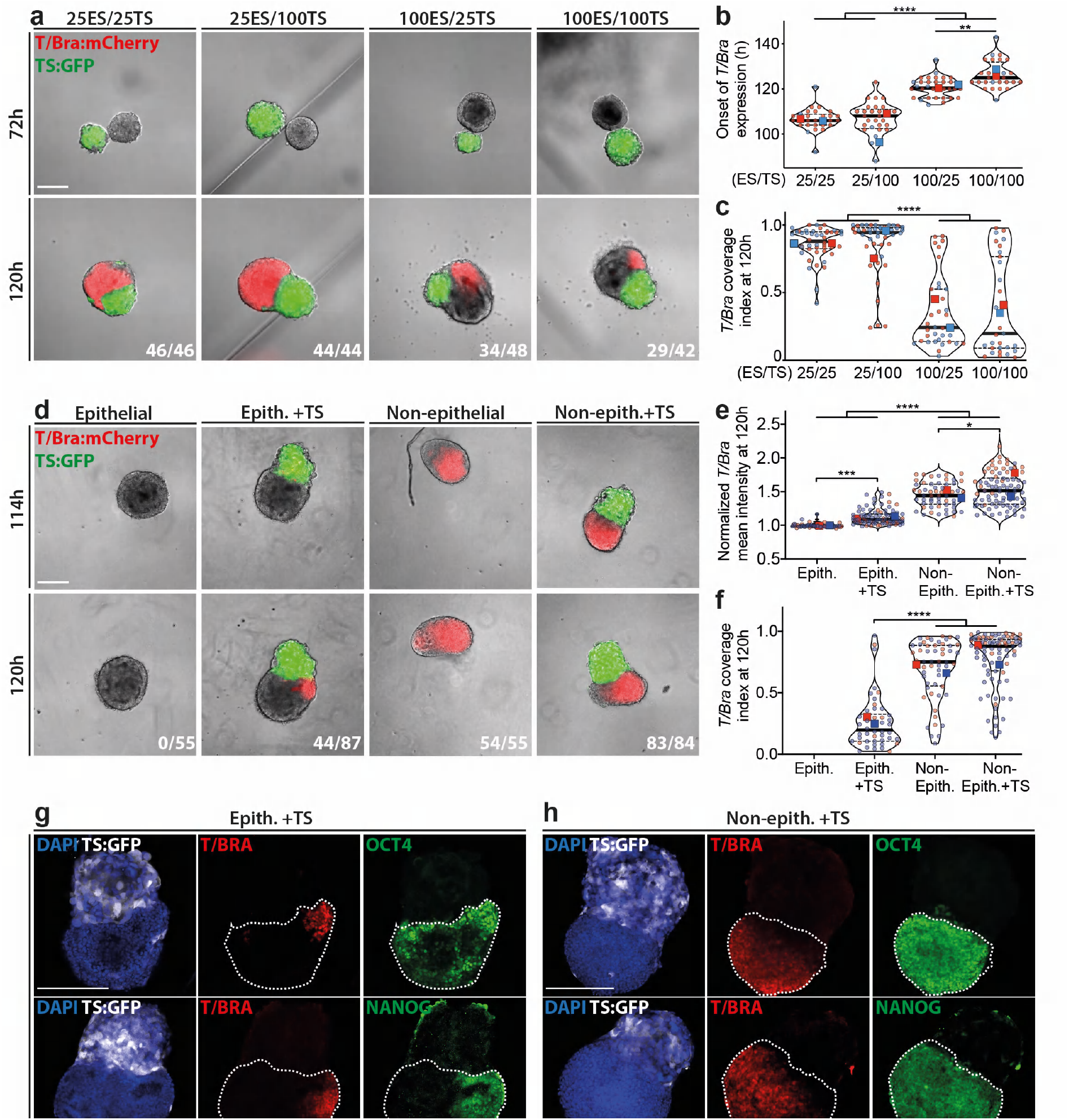
Effects of size and epithelial architecture of EPI aggregates on *T/Bra* expression dynamics in *EpiTS embryoids*. **a)** Representative images showing *T/Bra* expression dynamics in *EpiTS embryoids* formed from different starting cell numbers per well (ESC/TSC). **b)** Timelapse analysis between 78h and 149h with 2h interval showing the onset of *T/Bra* expression. For 25/25, 25/100, 100/25 and 100/100 conditions, total number of embryoids analyzed were 26, 27, 31 and 26, respectively. Data is collected from two biologically independent experiments. **c)** Coverage index of *T/Bra* expression calculated by division of *T/Bra*-positive area to EPI area. Data was collected from two biologically independent experiments. **d-f)** Representative images at 114h and 120h showing *T/Bra* expression **(d)**, background normalized mean intensity **(e)** and coverage **(f)** in epithelialized (+matrigel) or non-epithelialized (-matrigel) EPI aggregates, cultured in the presence or absence of TS aggregates at 120h. Data was collected from two biologically independent experiments. **g-h)** Representative confocal images at 120 hours showing *T/Bra, Oct4, Nanog, Otx2 and Sox2* expression in epithelialized **(g)** or non-epithelialized **(h)** embryoids. Nuclei were stained with DAPI. GFP-labeled TS cells were depicted in white. Dashed lines indicate EPI compartment. For all conditions in **(a-c)** and **(d-f)**, number of *T/Bra*:mCherry-positive embryoids over total number of embryoids analyzed are indicated at bottom right of (**a)** and **(d)**, respectively. Large symbols indicate mean values of each replicate. Black lines indicate median and quartiles. For all statistical analysis, one-way ANOVA followed by Tukey multiple comparison test was performed. Following P-value style was used: P****<0.0001, P***<0.0002, P**<0.0021, P*<0.0332. Scale bars: 200μm.

To test whether the appearance of *T/Bra* expression was dependent on the epithelial architecture of the aggregates, or the presence of an extraembryonic compartment, we generated epithelialized (with Matrigel) and non-epithelialized (w/o Matrigel) EPI aggregates, and cultured them in the presence or absence of TSC aggregates. Almost all non-epithelialized EPI aggregates displayed *T/Bra* expression by 120 h, independent of the extraembryonic compartment. In contrast, epithelialized EPI aggregates alone did not initiate *T/Bra* expression and required co-culture with TSC aggregates (**Fig. 2d,e**), similar to embryo explants cultured without extraembryonic ectoderm^25^. Moreover, the *T/Bra* expression domain was found to be more restricted in epithelialized *embryoids* (Epith.+TS) compared to non-epithelialized (Non-epith.+TS) ones (**Fig. 2f**).

To test whether *T/Bra* induction is specific to TSC aggregates, we co-cultured epithelialized EPI aggregates with aggregates composed of mouse embryonic fibroblasts (MEFs). MEF aggregates could occasionally induce *T/Bra* expression, albeit at much lower levels compared TSC aggregates (**Supplementary Fig. 4a,b**). This suggests that cellular interaction alone is not sufficient to induce *T/Bra* expression and that TSC aggregate-specific signaling may be necessary.

*In vivo*, signaling from the extraembryonic ectoderm is crucial for the initiation of gastrulation^26^. To better understand the signaling influence of the TSC compartment in our embryoids, we replaced the cellular aggregates with cell-adhesive hydrogel microbeads (composed of denatured collagen crosslinked to dextran) coated with various (diffusible) morphogens involved in early mouse development. When coupled to EPI aggregates at 72 h, plain beads or beads coated with Fgf2, Activin-A or Wnt3a could induce *T/Bra* expression, although at lower levels than TSC aggregates. Strikingly, Bmp4-coated beads reached the highest *T/Bra* expression level, outperforming the TSC aggregates (**Supplementary Fig. 4a,b**). Similarly, Bmp4 delivery increased WNT activity, even to higher levels compared to Wnt3a **(Supplementary Fig. 4a,c)**. Notably, we did not detect any significant change in TGF-β activity for the proteins tested **(Supplementary Fig. 4a,d)**. Transwell co-culture of epithelialized EPI aggregates with varying numbers of TSC aggregates did not induce *T/Bra* expression (**Supplementary Fig. 4c,d**), showing that physical interactions between the two cell compartment is crucial for symmetry breaking. Altogether, these results highlight a role of the epithelial architecture of EPI aggregates for *T/Bra* expression dynamics. EPI aggregates that are small (*i.e*., poorly epithelialized), or formed in the absence of Matrigel (*i.e*., not epithelialized), showed an extended *T/Bra* expression domain when coupled to TSC aggregates, suggesting uniform mesodermal differentiation, while in EPI aggregates the epithelium limits *T/Bra* expression to one side of the embryoids. Furthermore, TSC aggregate-specific signaling, likely via *Bmp4*, and physical EPI-TSC interaction are both important for the reproducible symmetry breaking and induction of *T/Bra* expression in epithelialized *EpiTS embryoids*.

### Cell fate assignments in *EpiTS embryoids*

Next, we performed immunostaining and confocal microscopy to characterize the tissues present in *EpiTS embryoids* at the onset of gastrulation. In the culture condition that yielded the most restricted induction of *T/Bra* expression, we observed that the lumen collapsed by 120 h and the *E-cadherin/Pdx-* positive epithelium disappeared in *T/Bra*-positive posterior domain, while it remained in the anterior domain where the cells remained in epithelial rosettes. Moreover, we detected a local downregulation of *laminin* where some *T-Bra*/*Snai1-*double positive cells emerged (**Supplementary Fig. 5a**), indicating basement membrane degradation and epithelial-to-mesenchymal transition (EMT) that precede migration from the primitive streak in the embryo^36,37^. Immunostaining for the pluripotency marker *Oct4* revealed uniform expression in all cells of the EPI compartment, partially overlapping with *T/Bra*-positive cells. The expression domain of *Nanog* was more restricted, encircling and co-localizing with *T/Bra* expression^3^. The epiblast marker *Otx2* was detected in a majority of the cells in the embryonic domain, excluding the *T/Bra*-positive cells. Notably, *Sox2* was solely expressed opposite and excluded from the *T/Bra* domain (**Fig. 2g**). We believe that the *Oct4+ T-Bra+ Nanog+* domain in these early *EpiTS embryoids* resembles the posterior end of the early to mid-streak embryo^27^. Conversely, the *Oct4+ Otx2+ Sox2+* domain marked the opposite end of the *embryoids*, suggesting maintenance of the anterior pluripotent epiblast. We did not detect *Sox17*-positive cells in *embryoids* that initiated *T/Bra* expression, but we could detect some *Dppa3*-positive cells (**Supplementary Fig. 5b**), likely marking primordial germ cell (PGC) progenitors^28^.

*Embryoids* formed from non-epithelialized EPI aggregates showed dispersed *T/Bra* expression and did not demonstrate *E-cadherin+* rosettes. Interestingly, *Pdx* expression was downregulated in the embryonic compartment. Similarly, *laminin* was completely absent, and the majority of cells demonstrated *Snai1* expression, suggesting an expanded primitive streak-like domain (**Supplementary Fig. 5c**). *T/Bra* expression mostly co-localized with *Oct4*/*Nanog*, however, *Otx2/Sox2* expression spanned a smaller domain compared to epithelialized *embryoids*, suggesting the loss of anterior neural progenitors at the expense of expanded posterior tissue (**Fig. 2h)**. Moreover, we detected a higher number of *Sox17*-positive endoderm-like cells and *Dppa3*-positive PGC progenitors (**Supplementary Fig. 5d)**, indicating an increased differentiation of posterior cell types.

Collectively, these results show that the epithelial architecture of the EPI aggregates directly influences the type of tissue that is created in *embryoids*. The presence of an epithelium favors limited *T/Bra* induction and a larger domain of anterior epiblasts that resemble E6.5 embryos. Conversely, in the absence of an epithelium in the initial EPI aggregate, the *embryoids* consist largely of posterior tissues and show premature differentiation towards the endoderm lineage.

### Gastrulation-like events are orchestrated by key early developmental signaling pathways

The onset of gastrulation is the output of asymmetric signaling across the post-implantation epiblast and is tightly regulated by WNT, TGF-β and BMP signaling pathways^29^. Embryos that lack functional β-catenin^30^ or cleavable Nodal^31^ fail to upregulate *T/Bra* and are unable to undergo gastrulation. Furthermore, embryos lacking both copies of *Bmp4* are arrested in development and do not express *T/Bra*^26^. The dependence of the symmetry breaking dynamics on the epithelial architecture of *EpiTS embryoids* led us to probe the underlying signaling pathways that regulate *T/Bra* expression by generating *EpiTS embryoids* from WNT (TLC:mCherry^32,33^) and TGF-β (AR8:mCherry^34^) reporter ESC lines and by using specific pathway inhibitors.

Time-lapse analysis showed that in non-epithelized *embryoids, T/Bra* expression begins as early as 96 h, in contrast to epithelized ones, which show a delayed initiation at 120 h (**Fig. 3a-b**, upper panels; **Fig. 3c**). In epithelialized *embryoids*, we could detect an increase of WNT activity from 72 to 96 h, followed by a reduction and restriction to the embryonic-extraembryonic interface by 120 h. In contrast, non-epithelial *embryoids* demonstrated a progressive increase in WNT activity that was detected throughout the EPI domain (**Fig. 3a-b,** middle panels; **Fig. 3d**). On the other hand, TGF-β signaling was continuously active in both conditions, but its expression domain became restricted over time. Of note, compared to epithelialized *embryoids*, non-epithelialized ones showed significantly higher TGF-β pathway activity from 72 to 120 h, (**Fig. 3a-b,** bottom panels; **Fig. 3e**). These results showed that in *EpiTS embryoids*, both WNT and TGF-β pathways are active and their dynamics depend on the epithelial architecture of the EPI aggregate. In epithelialized *embryoids*, WNT and TGF-β signaling preceded *T/Bra* induction and are co-localized by 120 h, resembling the initial primitive streak-like domain. The signaling polarization rendered the anterior of the *embryoids* low in WNT and TGF-β activity (**Fig. 3a-b**, white arrows), potentially preserving the potential for anterior neural progenitors^35,36^. In non-epithelialized *embryoids*, dispersed *T/Bra* expression was accompanied by elevated WNT and TGF-β activity, resembling an expanded posterior domain observed in Dkk1-/- and Cer1-/-;Lefty1-/- mutant embryos^37,38^.

**Figure 3:**
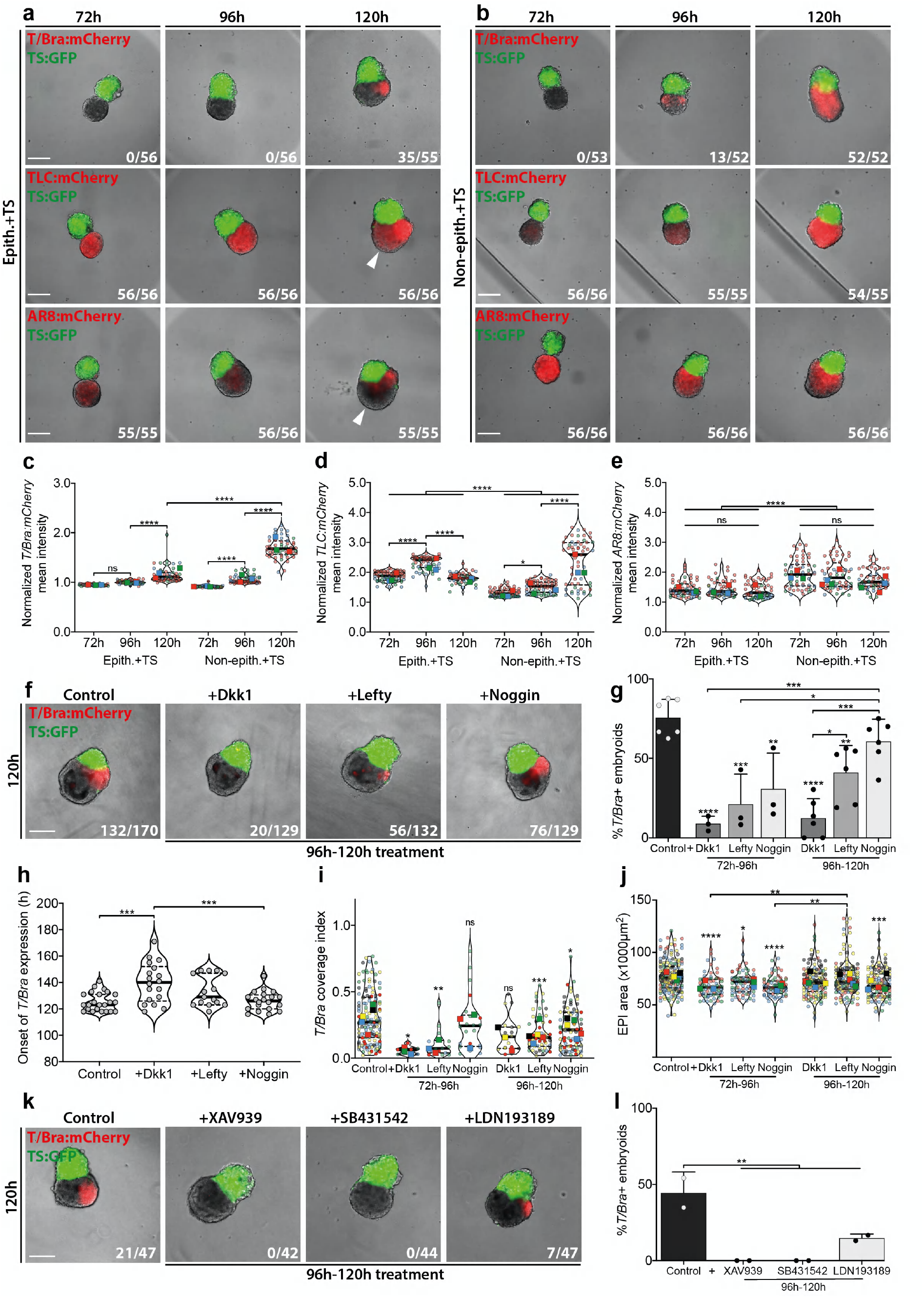
Roles of WNT, TGF-β and BMP pathways in *T/Bra* expression dynamics. **a-b)** Representative images showing *T/Bra:*mCherry (top), *TLC:*mCherry (middle) and *AR8*:mCherry (bottom) expression dynamics in epithelialized **(a)** and non-epithelialized **(b)** *EpiTS embryoids* between 72h to 120h. **c-e)** Background normalized mean intensity of *T/Bra:*mCherry **(c)**, *TLC:*mCherry **(d)** and *AR8*:mCherry **(e)** in epithelialized or non-epithelialized embryoids between 72h to 120h. Data was collected from three biologically independent experiments. **f)** Representative images at 120h showing *T/Bra:*mCherry expression in epithelialized embryoids treated with indicated inhibitors between 96h to 120h. **g)** Percentage of *T/Bra:*mCherry-positive epithelialized *embryoids* at 120h, treated with indicated inhibitors between 72h to 96h or 96h to 120h. Background offset was used to set a threshold for *T/Bra:*mCherry expression. **h)** Timelapse analysis between 78h and 192h with 2h interval showing the onset of *T/Bra* expression. For control, Dkk1, Lefty, Noggin conditions, total number of embryoids analyzed were 23, 18, 15, 20, respectively. Data is collected from a single experiment. **i-j)** Quantification of coverage index of *T/Bra:*mCherry expression **(i)** and EPI compartment area **(j)** at 120h in epithelialized embryoids treated with indicated inhibitors. **k)** Representative images showing *T/Bra:*mCherry expression in epithelialized embryoids treated with indicated small molecule inhibitors between 96h to 120h. **l)** Percentage of *T/Bra:*mCherry-positive epithelialized *embryoids* at 120h, treated with indicated inhibitors between 96h to 120h. For all conditions in **(a-e), (f-g,i-j)** and **(k-l)** number of mCherry-positive embryoids over total number of embryoids analyzed are indicated at bottom right of (**a-b)**, **(f)** and **(k)** respectively. Large symbols indicate mean values of each replicate. Black lines indicate median and quartiles. For all statistical analysis, one-way ANOVA followed by Tukey multiple comparison test was performed. For **(g)**, **(i)** and **(j)** significance was calculated in comparison to control. Following P-value style was used: P****<0.0001, P***<0.0002, P**<0.0021, P*<0.0332. Scale bars: 200μm.

To test whether gastrulation-like processes observed in epithelialized *EpiTS embryoids* involve mechanisms similar to those observed in the embryo, we inhibited the above pathways by adding Dkk1 (*i.e*., a WNT inhibitor), Lefty (TGF-β inhibitor) and Noggin (BMP inhibitor) between 72-96 h or 96-120 h, respectively. Inhibition of WNT and TGF-β pathways in both time windows greatly reduced the number of *embryoids* that upregulate *T/Bra*. However, BMP inhibition by Noggin between 72-96 h significantly lowered the percentage of *T/Bra*+ *embryoids* by 120 h (**Fig 3f,g)**. Dkk1-treated *EpiTS embryoids* exhibited a delayed *T/Bra* expression compared to non-treated or Noggin-treated ones (median ~140 h versus ~125 h) (**Fig 3h)**. Furthermore, we observed more restricted *T/Bra* expression in the presence of the inhibitors, which was more pronounced in the cases of Dkk1 and Lefty (**Fig. 3i**). Interestingly, early treatment with these inhibitors significantly reduced the total embryoid size, an effect that was more prominent when BMP signaling was inhibited (**Fig 3j)**. Treatment with the small molecule inhibitors XAV939 (*i.e*., a WNT inhibitor), SB431542 (TGF-β inhibitor) and LDN193189 (BMP inhibitor) resulted in a more pronounced reduction of *T/Bra* expression; SB431542 and XAV939 treatment led to a complete loss of *T/Bra* expression, while a small percentage of LDN193189-treated *EpiTS embryoids* still expressed *T/Bra*, yet with a domain that was spatially restricted (**Fig. 3k,l**). Altogether, these results demonstrate a crucial role of WNT and TGF-β signaling in the induction and level of *T/Bra* expression in *EpiTS embryoids* between 72 and 120 h. In contrast, the BMP pathway was shown to have an effect on the overall growth of *embryoids*, with a smaller effect on *T/Bra* expression between 96 to 120 h.

### Spontaneous axial morphogenesis of *EpiTS embryoids*

Previous embryoid models involving ESCs and TSCs in ETS-embryos^11^ did not recapitulate the axial patterning of the early embryo. In our hands, ETS-embryos could be generated at frequency (corresponding to multicellular aggregates comprising an embryonic and extraembryonic compartment) of ~6% (n=34/560) (**Supplementary Fig. 6a**). The ETS-embryos expressed epiblast markers *Oct4* and *Otx2* at 120 h (**Supplementary Fig. 6b**). To test whether post-gastrulation developmental stages could be achieved in ETS-embryos, we adopted our *EpiTS embryoid* culture approach by placing individual ETS-embryos in U-bottom 96-wells. *T/Bra* expression was found to be highly variable, with some tissues showing a dispersed and others a more localized expression by 120 h (**Supplementary Fig. 6c**). When cultured until 240 h, ETS-embryos with small embryonic-to-extraembryonic ratio were overgrown by TSC aggregates, and ETS-embryos that were small in size (< 200μm) did not grow further. Interestingly, we observed that ETS-embryos that upregulated *T/Bra* and had a larger embryonic compartment occasionally underwent axial elongation, suggesting that when grown in ECM- and serum-free culture conditions, these *embryoids* might reach post-gastrulation stages (**Supplementary Fig. 6d**).

When 120 hours old *EpiTS embryoids* were cultured in N2B27 medium with no additional growth factors, the embryonic compartment began to elongate (**Fig. 4a,b**). While epithelialized EPI aggregates alone remained spherical, non-epithelialized ones elongated efficiently, similar to classical gastruloids^39^ (**Supplementary Fig. 7a**). The EPI compartment in *EpiTS embryoids* grew significantly over time, while the size of the TSC aggregates did not change markedly (**Fig. 4c**), suggesting that growth of extraembryonic compartment is not required for the elongation of the EPI domain. To quantify the dynamics of elongation, we performed image analysis to determine the axial length and elongation index of the *embryoids* (**Supplementary Fig. 7b**). This analysis revealed that in both epithelialized and non-epithelialized *embryoids*, the axial length and the elongation index significantly increased from 120 to 168 h (**Fig. 4d, Supplementary Fig. 7c**). *EpiTS embryoids* formed from non-epithelialized (or smaller) EPI aggregates were significantly more elongated at 144 h (**Fig. 4d, Supplementary Fig. 7d,e**), suggesting a role for the size and epithelial architecture of the starting EPI aggregate on the elongation dynamics **(Supplementary Movies 5,6)**.

**Figure 4:**
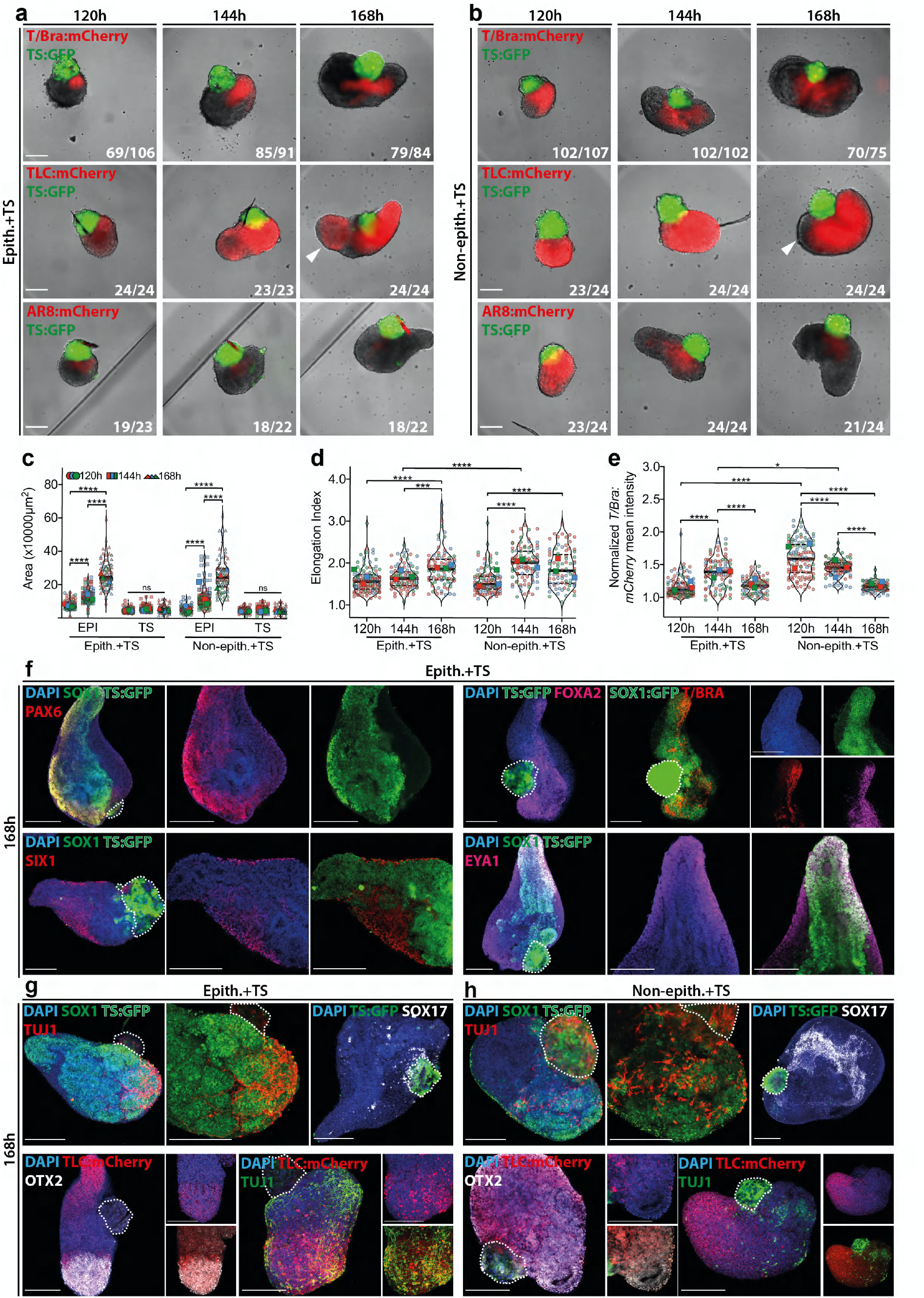
Axial morphogenesis in EpiTS embryoids. **a-b)** Representative images showing *T/Bra:*mCherry (top), *TLC:*mCherry (middle) and *AR8*:mCherry (bottom) expression dynamics in epithelialized **(a)** and non-epithelialized **(b)** *EpiTS embryoid* between 120h to 168h. **c-e)** Quantification of total EPI vs TS area **(c)**, elongation index **(d)** and background normalized *T/Bra:*mCherry mean intensity **(e)** in epithelialized or non-epithelialized embryoids between 120h to 168h. For elongation index quantification, TS subtraction was performed. Data was collected from three biologically independent experiments. **f)** Representative confocal images showing *Sox1, Pax6, T/Bra, Foxa2* (top panel) *Six1, Eya1* (bottom panel) immunostainings at 168h in epithelialized embryoids. **g-h)** Representative confocal images showing *Sox1, Tuj1, Sox17* (top panel) and *mCherry, Otx2, Tuj1* (bottom panel) immunostainings at 168h in epithelialized **(g)** or non-epithelialized **(h)** embryoids. Nuclei were stained with DAPI. TS cells were depicted with dashed line. For all conditions in **(a-e)**, number of mCherry-positive embryoids over total number of embryoids analyzed are indicated at bottom right of (**a-b)**. Large symbols indicate mean values of each replicate. Black lines indicate median and quartiles. For all statistical analysis, one-way ANOVA followed by Tukey multiple comparison test was performed. Following P-value style was used: P****<0.0001, P***<0.0002, P**<0.0021, P*<0.0332. Scale bars: 200μm.

In *embryoids* generated from epithelialized tissues, *T/Bra* expression peaked around 144 h and gradually declined until 168 h, when it became restricted to the elongating tip and then trailed behind in a short stripe of cells. In contrast, non-epithelialized *embryoids* showed a constant decrease in the intensity and size of the *T/Bra* domain with increasing elongation (**Fig. 4e, Supplementary Fig. 7f**). WNT signaling was active in almost all *embryoids* and marked the *T/Bra-*expressing tip with no significant change in activity levels until 168 h (**Supplementary Fig. 7g,h**). In epithelialized *embryoids*, WNT activity reappeared at the anterior domain (**Fig. 4a,b,** middle panels, white arrows), resembling the TCF/LEF reporter activity in the tail and midbrain/hindbrain of E8.5 embryos^33,40^. Overall, TGF-β pathway activity was found to be lower and in close proximity to the extraembryonic compartment, excluded from the elongating tip (**Fig. 4a,b,** bottom panels; **Supplementary Fig. 7i,j**).

### Signaling and axial patterning in *EpiTS embryoids*

To better understand the underlying mechanism of axial morphogenesis in our *embryoids*, we investigated the roles of the WNT, TGF-β and BMP pathways by inhibiting their activity between 96 and 120 h. In general, treatment with inhibitors resulted in significant size reduction by 168 h (**Supplementary Fig. 8a**). Dkk1 and Lefty led to a reduction of *T/Bra-*positive *embryoids*, compared to Noggin. *Embryoids* treated with Lefty exhibited small clumps of *T/Bra*-positive cells, while Dkk-treated ones showed a more scattered expression throughout the body of the structure (**Supplementary Fig. 8b**). In Noggin-treated *embryoids*, *T/Bra* expression was retained within a larger domain compared to untreated or Dkk1/Lefty-treated ones (**Supplementary Fig. 8c**). Notably, the elongation frequency was found to be significantly higher in the presence of Noggin (**Supplementary Fig. 8d, Supplementary Movies 7-10**). It is conceivable that excessive BMP signaling from the extraembryonic ectoderm could impair axial elongation, and treatment with Noggin could mitigate this effect, consistent with the antagonistic role of BMP4 on axial elongation of gastruloids^18^. In these *embryoids, T/Bra*-positive cells were often organized into long stripes that co-expressed *FoxA2*, suggestive of a notochord or an anterior mesendoderm (AME)-like structure (**Supplementary Fig. 8e**). In support of this, we detected higher expression levels of *T/Bra, Shh, FoxA2* and *Gsc* with Noggin-treated compared to untreated *embryoids* (**Supplementary Fig. 8f**), in line with previous reports defining a role for Noggin in proper notochord formation^41^.

At 168 h, epithelialized *embryoids* acquired an expanded neural identity, as evidenced by *Sox1* expression spanning from the elongated tip towards the opposite end. Interestingly, we could detect polarized expression of *Pax6* and *T-Bra/Foxa2*, all partially overlapping with *Sox1* expression, suggesting a dorsal-ventral organization similar to the developing neural tube^42^. Furthermore, the neural tissue was flanked by *Six1+ Eya1+* paraxial mesoderm derivatives (**Fig. 4f**), suggesting an organization across a medio-lateral axis. We detected numerous *Tuj1*-positive neurons within *Sox1*-positive rosettes located on the opposite end of the elongated tip. The rosettes and neurons showed high WNT activity co-localized with *Otx2* expression, indicating establishment of an anterior-posterior axis to include a ‘brain-like’ tissue. Moreover, only few *Sox17*-positive cells were found scattered in the vicinity of the extraembryonic domain, suggesting poor endoderm differentiation (**Fig. 4g**). In contrast, non-epithelialized *embryoids* showed a smaller domain of *Sox1* expression comprising comparable numbers of *Tuj1*-positive neurons with relatively short axonal projections. Compared to epithelialized *embryoids*, the *Otx2* expression domain was more restricted and showed low levels of WNT activity. *Tuj1*-positive neurons were also located away from the WNT-positive elongated tip. Interestingly, the expression domain of *Sox17* in these *embryoids* was much larger and located at the anterior end of the *embryoids* co-localizing with *Foxa2* (**Fig. 4h, Supplementary Fig. 8g**), similar to endoderm tissue seen in gastruloids^6^. Collectively, these data demonstrate the capacity of *EpiTS embryoids* to spontaneously undergo axial morphogenesis to establish multi-axial patterning. Importantly, the presence or absence of an epithelium in the starting EPI aggregate directly influences the elongation dynamics as well as the proportions of the tissue types that are generated.

### Single cell RNA-sequencing reveals distinct differentiation trajectories in *EpiTS embryoids*

To shed light on the cell type composition and diversity of *EpiTS embryoids*, we performed single cell RNA sequencing (scRNA-seq) analysis between 120 and 192 h (**Fig. 5**). We could identify that from 120 h onwards, cells mapping to an epiblast-state gradually differentiated into ectoderm or mesendoderm lineages (**Fig. 5a,** germ layers). Primordial germ cells and extraembryonic tissue had a distinct transcriptional identity that segregated away from the germ layers observed in our *embryoids*. RNA velocity on extraembryonic cells showed no clear temporal differentiation trajectory; however, we identified distinct cell populations that suggested a compartmentalization within the extraembryonic tissue (**Fig. 5a,** extraembryonic). At 120 h, the majority of the extraembryonic tissue had chorion identity (*Cdx2, Elf5, Eomes, Esrrb*) and expressed *Bmp4*, supporting involvement of BMP signaling for the initiation of gastrulation-like events in our *embryoids*. Interestingly, we detected *Furin* and *Pcsk6* expression that could suggest the processing of embryonic Nodal by the extraembryonic tissue^43^. We further identified clusters of cells that map to ectoplacental cone (*Fgfbp1, Ascl2*) or labyrinth progenitors (*Gjb2, Dlx3, Ovol2*) that demonstrate multipotentiality of chorionic ectoderm. At later stages of culture, we detected trophoblast giant cells (*Hand1, Ctsq, Prl2c2*) and a few spongiotrophoblast cells (*Tpbpa, Flt1*) (**Fig. 5b,** extraembryonic markers**, Supplementary Fig. 9a-d**), in line with previous *in vitro* TSC differentiation data^44^.

**Figure 5:**
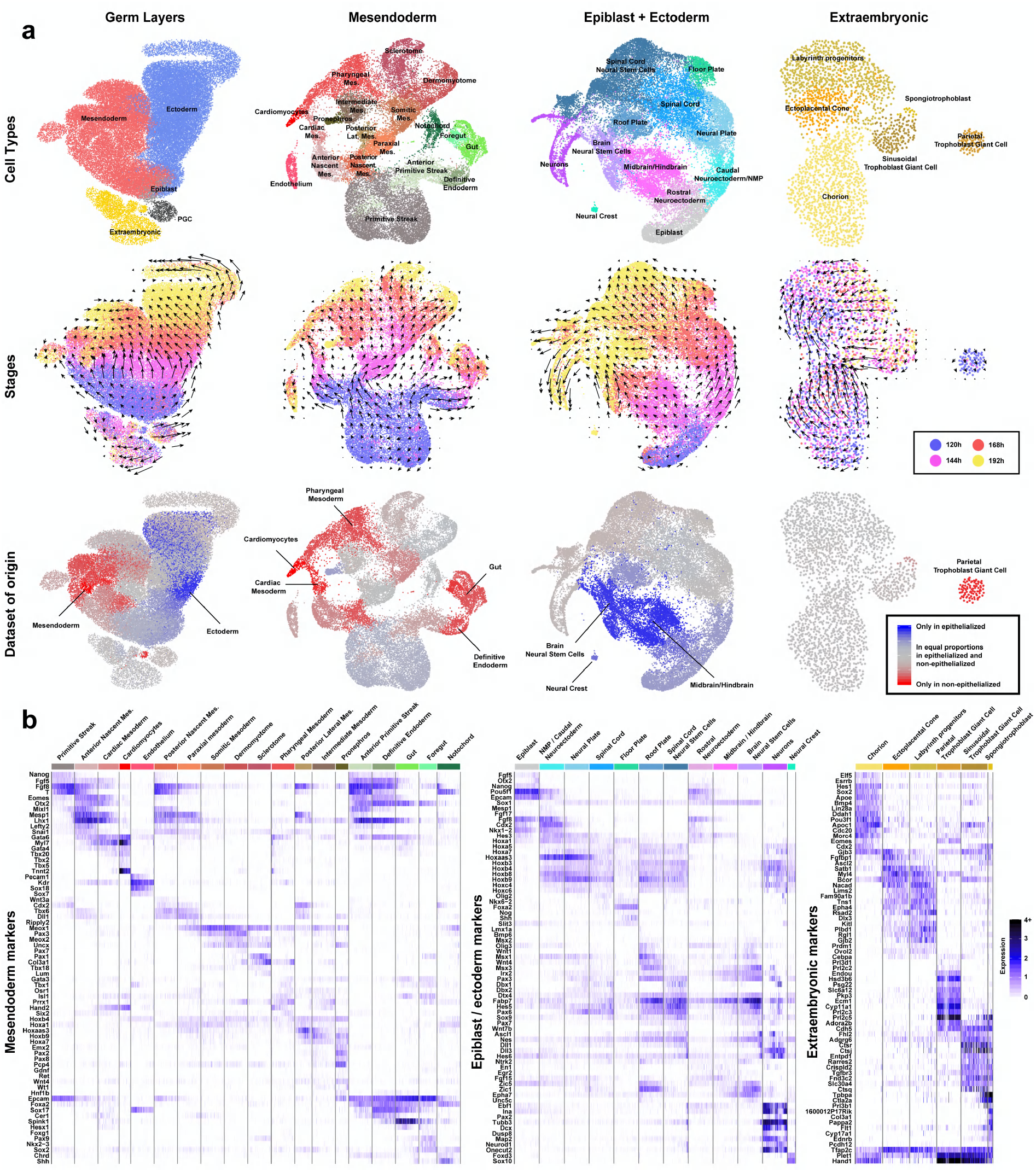
Comparative single cell RNA sequencing analysis of EpiTS embryoids. **a)** UMAP plots showing cell types, RNA velocity overlayed stages and datasets of origins in germ layers, mesendoderm, epiblast+ectoderm and extraembryonic clusters. **b)** Heatmap plot of selected marker genes from mesendoderm, epiblast+ectoderm and extraembryonic clusters. *PGC=Primordialgerm cells, Lat.=Lateral, Mes.=Mesoderm*

A close up on the mesendoderm cluster revealed complex differentiation trajectories (**Fig. 5a,** mesendoderm). At 120 h, the majority of cells had a primitive streak identity (*T, Mixl1, Wnt3*), that progressively branched to give rise to endoderm (*Sox17, Spink1*) and notochord (*Noto, T, Shh*) tissues via an anterior primitive streak-like population (*Foxa2, Gsc*). Cells mapping to the nascent mesoderm (*Mesp1, Hoxb1*) diverged into two distinct trajectories: the anterior portion gave rise to cardiac mesoderm (*Myl7, Gata4, Hand2*) and cardiomyocytes (*Ttn, Nkx2-5, Mef2c*)^7^, while the posterior nascent mesoderm cells differentiated towards paraxial mesoderm (*Msgn1, Tbx6, Meox1*). The paraxial mesoderm further diverged into somitic (*Meox2, Pax3, Uncx*), posterior lateral mesoderm (*Msx2, Foxf1, Hoxc6*) or intermediate mesoderm (*Osr1, Gata3, Wt1*) tissues. From 168 to 192 h of culture, the abovementioned mesoderm populations gave rise to more differentiated pharyngeal mesoderm (*Prrx1, Isl1, Tbx1*), sclerotome (*Pax1*) and dermomyotome (*Dmrt2*) cells (**Fig. 5b,** mesendoderm markers**, Supplementary Fig. 10a,b**).

A detailed RNA velocity analysis of the ectoderm cluster revealed that between 120 and 144 h, epiblast cells (*Oct4, Nodal*) gave rise to spinal cord tissue (*Sox1, Pax6, Hoxb7/8/9*) (**Fig. 5a,** epiblast+ectoderm). We noted a striking compartmentalization among the cells of the spinal cord, with differentiation towards dorsal (*Wnt1, Dbx2, Pax7*) and ventral (*Olig2, Foxa2, Shh*) cells of the neural tube. On a second trajectory, epiblast cells differentiated to generate midbrain/hindbrain tissue. The lower expression level of midbrain markers (*En1/2, Pax5*) and higher expression level of hindbrain markers (*Egr2, Irx2, Epha7, Fst*) could suggest that the ‘brain-like’ tissue generated in the *embryoids* was primarily of hindbrain origin. In support of this, we detected *Hoxa2, Hoxb1/2* and *Cyp26* expression in the hindbrain-like tissue, while more posterior *Hoxa3/b3* and *Hoxb4* were excluded from this region (**Fig. 5b,** epiblast+ectoderm markers; **Supplementary Fig. 11a,b**), suggesting that the hindbrain tissue in our *embryoids* corresponds to rhombomeres 1-6^45^. At later stages, we detected neural stem cells (*Pou3f3, Ascl1, Nes*) that either had midbrain/hindbrain (*Zic1/5, Msx3*) or spinal cord (*Hoxa4/b4*) identities. These neural stem cells generated dorsal interneurons (*Lhx1/5, Pou4f1*) and ventral motor neurons (*Isl1/2, Nkx6-1 Phox2a/b*) at both spinal cord (*Hoxb7/8/9*) and hindbrain (*Lhx2/9, Barhl1/2*) levels (**Supplementary Fig. 12a-d**), revealing a remarkable recapitulation of the cell type diversity similar to the *in vivo* situation^46^.

We then used the scRNA-seq dataset to systematically assess the effect of the initial epithelial architecture of *embryoids* on the resulting cell type diversity. A comparative analysis of epithelialized and non-epithelialized *embryoids* showed that several tissue types, including cardiac mesoderm, definitive endoderm and pharyngeal mesoderm derivatives, were almost exclusively found in non-epithelialized *embryoids* (**Fig. 5e, Supplementary Fig. 13a,b**). Moreover, the neural tissues in non-epithelialized *embryoids* were shown to be composed of spinal cord tissue. Conversely, midbrain/hindbrain tissue, brain-specific neural stem cells and their respective neurons were only detected in epithelialized *embryoids* (**Supplementary Fig. 12e**). These results demonstrate that the epithelial architecture of initial EPI aggregates has a direct impact on embryonic developmental trajectories and promotes neural differentiation in *embryoids*.

## Discussion

We have developed a novel bioengineered embryoid culture system for the study of early mouse postimplantation development that combines aggregates of ESCs and TSCs. Unlike previous studies^11,12,47^, this culture method allows simple independent modulation of the embryonic and extraembryonic compartments and makes our system both experimentally tractable and efficient for quantitative and mechanistic studies.

The controlled interaction between EPI and TSC aggregates leads to *EpiTS embryoids* that resemble the early mouse embryo. Specifically, EPI aggregates that have an apicobasally polarized epithelium develop a polarity with high *T/Bra* and low *Sox2/Otx2* expression on one end, and no *T/Bra* and high *Sox2/Otx2* expression on the rest of the aggregate. Furthermore, at this time the lumen of the epithelium collapses and while the *T/Bra* expressing cells appear to undergo an EMT, cells on the opposite side remain organized in epithelial rosettes. This organization resembles the anterior-posterior axis of the E6.5 embryo and strictly depends on the initial size of the EPI aggregate. We identify BMP signaling from the extraembryonic compartment, via WNT/TGF-β pathways, as the key regulator of polarized *T/Bra* expression in *EpiTS embryoids*.

A significant finding of our study is the effect that an epithelium has on *EpiTS embryoid* development. Under the same culture conditions, *T/Bra* expression is much more restricted in epithelialized than in non-epithelialized EPI aggregates. This suggest that the formation of an epithelium raises the threshold for WNT and TGF-β signal response such that, after the onset of *T/Bra* expression at one end, the probability that a second event happens in the aggregate end is low and this contributes to the restricted patterning of the EPI. The culture conditions we use here sensitize the EPI compartment to *T/Bra* expression which, in non-epithelialized aggregates, initially spreads throughout the aggregate. However, in the original gastruloid protocol, ESC aggregates undergo spontaneous symmetry breaking in the absence of any extraembryonic signals, localized to one end of the aggregate. Altogether, these observations suggest that one role of the extraembryonic ectoderm is to bias the spontaneous symmetry breaking event and that the epithelial organization of the EPI contributes to this by raising the threshold for signal response. The mechanism for this is unclear at the moment but the junctional organization of signaling receptors in the epiblast^48,49^ is likely to play a role. The situation we observe in *EpiTS embryoid* phenocopies aspects of the activity of the AVE where BMP, TGF-β and WNT signaling inhibitors secreted from the Anterior Visceral Endoderm (AVE) inhibit the spread of *T/Bra* expression, contributing to the anterior-posterior patterning of the embryo, suggesting that the epithelial organization of the epiblast contributes to this effect.

Long-term culture of *EpiTS embryoids* in growth factor-free medium resulted in axial elongation (originating from the *T/Bra* expressing pole) and subsequent multi-axial patterning. The extent of this process was dependent on the proportions of cell types generated as a function of the initial tissue architecture of the EPI aggregates. For example, we detected cardiac mesoderm and definitive endoderm derivatives almost exclusively in non-epithelialized *embryoids*, resembling post-occipital patterning in *gastruloids*^6^. In both cases the EPI compartment displayed expanded WNT and TGF-β signaling at early stages of culture, and this is likely to cause the observed loss of anterior neural progenitors. In contrast, epithelialized *embryoids*, although also exhibiting *T/Bra* expression initially, later are depleted for mesodermal derivatives and display cell fates associated with midbrain/hindbrain and, in some instances, spinal cord levels, supporting the importance of an epithelium for the formation of brain tissues^50,51^. The organization of epithelialized *embryoids* can be construed as an ‘early embryo’ in which the primitive streak is initiated but immediately truncated, as they lacked most of the mesendodermal derivatives. This could suggest that neural induction has taken place in these embryoids. In support of this, late *EpiTS embryoids* had *T/Bra*+*Foxa2*+ tissue extending along the embryoids suggesting the existence of a notochord/anterior mesendoderm (AME)-like structure which in the embryo is responsible for the induction of the midbrain/hindbrain^52,53^.

A surprising feature of the emergence of brain structures in the *EpiTS embryoids* is that it occurs in the absence of an AVE which, in the embryo, is thought to be needed to preserve neural potential in the epiblast^23^. An earlier study has shown that gastruloids derived from a combination of XEN cells and ESCs trigger an epithelium^12^ and develop midbrain/hindbrain tissue^52^. In the embryoids that lack extraembryonic endoderm cells, we do not observe any forebrain tissue, supporting this possibility. This could potentially be overcome by WNT inhibition at later stages of the culture, since Dkk1 secreted from the prechordal plate is known to be instructive for forebrain formation^53^. Together with our results, these observations suggest that the AVE might be involved in maintaining an intact epithelium in the anterior region of the epiblast^54^, and that this might be sufficient to preserve the neural potential which is later developed by the AME during neural induction.

Our experiments show how bioengineering-inspired approaches can be exploited to standardize complex embryoid models and allow for the modeling of embryonic-extraembryonic cell/tissue interactions involved in the generation of precise patterns during development. *EpiTS embryoids* should serve as a canvas to generate developmentally relevant organ-specific structures of either neural or mesendoderm origin, through an *in vivo*-like gastrulation process. Traditional approaches to identify tissue origins in the embryo have relied largely on genetic manipulations which are labor intensive. *EpiTS embryoids* thus provide an alternative approach to decipher the roles of mechanics and gene expression, which is difficult to perform *in vivo*. Ultimately, the adoption of our approach to generate human *embryoids* holds significant potential to shed light on early human development as experimenting with early human embryos is challenging.

## Acknowledgements

We thank G. Rossi and S. Vianello for feedback on the manuscript. We thank Lorenzo Mattolini for help with cell culture and imaging as well as contributions to image analysis. We thank all members of the Lutolf laboratory for sharing materials and insightful discussions. We thank E. Friman and D. Suter for providing the SBr (*Sox1*:GFP;*T/Bra*:mCherry) ESC line and C. Schröter for providing the TS:GFP cell line. We thank R. Guiet and O. Burri from the Bioimaging and Optics Platform (BIOP) at EPFL for writing the codes for image analyses. This work was funded by a Sinergia grant (CRSII5_189956) from the Swiss National Science Foundation, the National Center of Competence in Research (NCCR) Bio-Inspired Materials, and EPFL.

## Contributions

M.U.G. and M.P.L. conceived the study, designed experiments, analyzed data and wrote the manuscript. M.U.G. performed the experiments. N.B.^1^ analyzed single cell RNA sequencing data. S.H. and N.B.^1,4^ developed and fabricated microwell arrays. B.M. performed experiments on testing cytoskeletal inhibitors on epithelium formation and helped with image analysis. A.M.A. contributed to study and experimental design, data analysis and manuscript writing.

## Competing Financial Interest

The Ecole Polytechnique Fédérale de Lausanne has filed for patent protection on the microwell array technology applied herein, and S.H., N.B. and M.P.L. are named as inventors on those patents; S.H., N.B. and M.P.L. are shareholder in SUN bioscience SA, which is commercializing those patents.

## Methods

### Cell culture

Mouse embryonic stem cells (SBr line^55^) were cultured at 37°C in 5% CO2 in medium composed of DMEM+Glutamax (#61965-026), 10% ES cell-qualified FBS (#16141-079), 1mM sodium pyruvate (#11360-070), 1x MEM non-essential aminoacids (#11140-035), 0.1mM 2-mercaptoethanol (#31350-010) and 1000u/ml Pen/Strep (#15140-122) supplemented with 3μm GSK3i (#361559), 2μm MEKi (#S1036) and 0.1μg/ml LIF (in house preparation). Cells were routinely passaged every 2-3 days by seeding 8000-9000 cells/cm^2^ and every 20 passages a fresh vial was thawed. Cells were tested and confirmed free of mycoplasma. Mouse trophoblast stem cells (TS:GFP line^45^) were cultured at 37°C in 5% CO2 in TS medium composed of RPMI 1640+Glutamax (#61870-010), 20% ES cell-qualified FBS (#16141-079), 1mM sodium pyruvate (#11360-070), 0.1mM 2-mercaptoethanol (#31350-010) and 1000u/ml Pen/Strep (#15140-122). TS medium was conditioned on irradiated MEFs for 3 days and stored at −20°C. This was repeated three times for one batch of irradiated MEFs. Aliquots of TS conditioned medium (TSCM) were thawed and mixed 3:1 with fresh TS medium before cell passaging. 50ng/ml Fgf4 (#100-31) and 1μg/ml Heparin (#H3149) were added to make final TS medium. TS cells were routinely passaged every 2-3 days by seeding 5000-6000 cells/cm^2^ and every 20 passages a fresh vial was thawed. Cells were tested and confirmed free of mycoplasma.

### Preparing EPI and TS differentiation medium

N2B27 medium was prepared by 1:1 mixing of DMEM/F12+Glutamax (#31331-028) and Neurobasal (#21103-049) with the addition of 0.5x N2 supplement (#17502001), 0.5x B27 supplement (#17504001), 0.5x Glutamax (#35050-038), 1mM sodium pyruvate (#11360-070), 1x MEM nonessential aminocacids (#11140-035), 0.1mM 2-mercaptoethanol (#31350-010) and 1000u/ml Pen/Strep (#15140-122). 12ng/ml Fgf2 (#PMG0035), 20ng/ml Activin-A (#338-AC) and 1% KSR (#10828-010) were added to make final EPI differentiation medium (EPIdiff). TS differentiation medium (TSdiff) was prepared by 1:1 mixing of N2B27 and TS medium supplemented with 25ng/ml Fgf4 (#100-31) and 500ng/ml Heparin (#H3149).

### Preparing EPI and TSC Aggregates on PEG microwells

Poly(ethylene glycol) (PEG) microwells with 400μm well diameter (121 wells per array) were prepared on 24-well plates as previously described^56^. Microwells were equilibrated with 50μl of either EPIdiff (for ES cells) or TSdiff (for TS cells) for at least 30 minutes at 37°C. Mouse ES and TS cells and were dissociated to single cells with Accutase (#A11105-01) or TrypLE (#12605-028), respectively. Cells were then centrifuged at 1000 rpm for 5 minutes and washed twice with 10 ml PBS at 4°C. Cells were resuspended in cold EPIdiff (for ES cells) or TSdiff (for TS cells) and suspension of 484.000 cells/ml was prepared. 35μl of the suspension was added dropwise on microwell arrays to have 100-150 cells/well. Seeding was done at 37°C for 15 minutes. Growth factor reduced Matrigel (#356231) was diluted in cold EPIdiff or TSdiff to 3% or 2% (v/v), respectively. The medium was vortexed and 1ml was slowly added from the side of the well, avoiding direct addition from the top of the microwell arrays. Plates were kept at 37°C in 5% CO2 for at least 72 hours before further processing.

### Forming EpiTS embryoids

At 72-75 hours of culture, aggregates on microwell arrays were flushed out and transferred to non-tissue culture treated 10cm plates in 10ml warm N2B27 medium. Single EPI and TSC aggregates were picked in 10μl and transferred to low adherent U-bottom 96 well plates (#COR-7007). 170μl of N2B27 medium was added on top. At 96 and 120 hours, 150μl of medium was replaced with fresh N2B27 and *EpiTS embryoids* were kept until 168h. Protein inhibitors Lefty (#746-LF-025), Noggin (in house preparation), Dkk1 (#5897-DK-010) were added at indicated timepoints at 200ng/ml final concentration. Small molecule inhibitors; SB431542 (#S4317), XAV939 (#S1180), LDN193189 (#SML0559) were added at indicated timepoints at 10μm, 10μm and 1μm final concentrations, respectively.

### Testing effect of cytoskeletal inhibitors on EPI aggregate epithelialization

EPI aggregates were formed in the presence of Matrigel with following inhibitors until 72h: 1μm LPA(#L7260), 10μm Y27632 (#72302), 10μm Blebbistatin (#72402), 1μm ML-7 (#4310), 5mm Calyculin-A (#9902S), 100nm Cytochalasin-D (#250255), 1μm Latrunculin-A (#L5163), 100nm Jasplakinolide (#J4580), 50μm CK-666 (#SML0006), 25μm BpV (#S8651).

### Culturing of EPI aggregates with protein coated beads or on transwells

For bead experiments, Cytodex 3 microcarriers (#17048501) were resuspended in HBSS (Ca++, Mg++) at 40,000 beads/ml concentration. 600 beads were transferred in 0.5ml Eppendorf tubes and Fgf2 (#PMG0035), Activin-A (#338-AC), Wnt3a (in house preparation) or Bmp4 (#120-05ET-100) was added at 25μg/ml final concentration. Beads were incubated at 4°C overnight. Next day, beads were washed repeatedly to reach 1:250000 dilution in N2B27 and transferred in 10μl to low adherent U-bottom 96 well plates together with epithelialized EPI aggregates.

For transwell experiments, EPI aggregates were placed in 190μl to low adherent U-bottom 96 well plates and 1, 3 or 5 TSC aggregates were placed in transwells on top (#CLS3374). 40μl medium was added to cover TSC aggregates. At 96h, 180μl of medium was replaced with fresh N2B27.

### Immunostaining and confocal microscopy

*EpiTS embryoids* were washed with PBS and fixed with 4% PFA for 2 hours at 4°C. PFA was removed by three serial washes of 20 minutes at room temperature. Blocking was performed in blocking solution (PBS+10%FBS+0.3% Triton-X) for 1 hour at room temperature. Primary antibodies (see supplementary table 1) were incubated for at least 24 hours at 4°C in blocking solution. Next day, primary antibodies were removed by three serial washes of 20 minutes at room temperature. Secondary antibodies were incubated for 24 hours and next day embryoids were washed and mounted on glass slides in mounting medium. Confocal images were taken using an LSM700 inverted (Zeiss) with EC Plan-Neofluar 10x/0.30 or Plan-Apochromat 20x/0.80 air objectives.

### Image analysis

All images were processed using algorithms developed in Image J (version 2.0.0-rc-69/1.52n). Brightfield, GFP (for TS cells), and mCherry (for *T/Bra*) channels were used as input. Thresholding and segmentation was performed sequentially for each channel. Area of EPI domain was calculated by subtraction of the area of the object identified in GFP channel from the area of the object identified in brightfield channel. For the calculation of the anterior EPI domain, area of the object identified in mCherry channel was subtracted from the EPI area. *T/*Bra coverage index was calculated by dividing area of the object identified in mCherry channel to the EPI area. For morphology measurements, bright-field images were thresholded and segmented. Maximum inscribed circle function was used to fit circles in the identified object. Axial length was determined by connecting centers of the fit circles. Elongation index was calculated by dividing axial length to the diameter of the maximum inscribed circle. All image analysis codes are available upon request.

### RNA isolation and qPCR

RNA was extracted with the RNeasy Micro kit (QIAGEN), according to manufacturer’s instructions and quantified with a spectrophotometer. 1 μg of RNA was reverse transcribed with the iScript cDNA Supermix (#1708890). cDNA was diluted 1:5 and amplified by using Power SYBR Green PCR Master Mix. qPCR was run with a 7900HT Fast PCR machine (#4329001), using Power SYBR Green PCR Master Mix (Applied Biosystems), with an annealing temperature of 60°C. Gene expression was normalized on Gapdh expression. Relative fold expression was calculated with the 2-ΔΔCT method. Primers used are indicated in the Supplementary Table 2.

### Bulk RNA-seq

*EpiTS embryoids* were lysed with 200 μl trizol, followed by addition of 70 μl of chloroform to trigger phase separation and then the aqueous phase was collected. The extraction process was repeated a second time, and an equal volume of isopropanol was added to precipitate the RNA, which was collected by centrifugation at 20000 g for 30 min. The pellet was washed with 15 mM sodium acetate in aqueous 70% ethanol, followed by salt-free 70% ethanol, before picking up in RNAse free water. RNA quantity and quality were assessed on nanodrop, qubit, and Agilent TapeStation 4200 profiling, and showed absorbance ratios 260/280 of 1.85 ± 0.12 and RNA integrity numbers (RIN) of 9.9 ± 0.2 (average ± SD), supporting good purity and absence of degradation. TruSeq stranded mRNA LT libraries were prepared according to Illumina protocol 15031047 Rev. E, starting from 300 ng of RNA, quantified by qubit DNA HS, profiled on TapeStation 4200, and sequenced on an Illumina HiSeq 4000 at a targeted depth of 36 Mreads/sample and paired-end read length of 81,8i,8i,81. The reads were trimmed for adapters with bcl2fastq v2.20.0, aligned to the mouse genome mm10 with STAR 2.7.0e, and a count matrix was assembled using the cellranger v4.0 curation of ENSEMBL annotations. In the manuscript, “Gene expression” refers to natural logarithm of counts per million for bulk RNA-seq data. The data was collected from 4 independent experiments.

### Single-cell RNA-seq

Single cell RNA-seq of *EpiTS embryoids* was performed with 10x single cell 3’ gene expression reagent kit chemistry v3.1, according to the protocol CG000204 Rev D. Library QC was performed on an Agilent TapeStation 4200, and sequencing was performed on Illumina HiSeq 4000, with a paired-end read length of 28, 8i, 110, targeting approximately 3000 cells and 300 Mreads per library (which included two replicates per condition, 4 timepoints, epithelialized and non-epithelialized embryoids, for a total of 16 libraries). Recovered cells were 3300 ± 900, and reads per cell after UMI collapse of 29 000 ± 4500 (average ± SD). The reads were trimmed for adapters and polyA with cutadapt^57^ v2.1, and aligned to mm10 with cellranger 3.1. The count matrices were imported in R v3.6 with Seurat^58^ v3.1. Live cells were selected based on detection of more than 2000 genes, and mitochondrial gene content of 1.5 to 15%, and counts were normalized as natural logarithm of counts normalized to 10k per cell (referred to as “Gene expression” in the manuscript). A mini-analysis including dimensionality reduction by PCA based on centered and scaled most variable genes and cell type annotation transfer from an *in vivo* atlas^59^ with scmap^60^ was performed on individual datasets for the needs of doublet removal with DoubletFinder v2.0.3^61^. After data filtering, all datasets were merged, variable genes, scaling and PCA repeated, and batch correction was then done with Harmony v1^62^. A first Louvain clustering yielded clusters that were assigned to germ layers. UMAPs^63^ were computed with uwot v0.1.5, first with 100 cells per cluster and spectral initialization, and then with all cells, initialized on the previous cluster centroids plus noise. The data was then split by germ layers, and a refined Louvain clustering and UMAP view was computed for each subset. Louvain clusters were finally assigned to cell types based on canonical markers. Custom code was used for visualizing cell type proportions vs day and/or epithelialization, normalizing for heterogeneous dataset sizes to avoid bias. A further closeup was similarly performed on the neurons of the epiblast+ectoderm subset. Splicing data was estimated with velocyto^64^ v0.17, and RNA-velocity was computed with scVelo v0.2.1^65^ in deterministic mode, using python v3.7.2 and scanpy v1.5^66^.

## Notes

### Competing Interest Statement

The EPFL has filed for patent protection on the microwell array technology applied herein, and S.H., N.B. and M.P.L. are named as inventors on those patents; S.H., N.B. and M.P.L. are shareholder in SUN bioscience SA, which is commercializing those patents.

## References

1. Fu, J., Warmflash, A. & Lutolf, M. P. Stem-cell-based embryo models for fundamental research and translation. Nature Materials 20, 18–13 (2020).

2. Warmflash, A., Sorre, B., Etoc, F., Siggia, E. D. & Brivanlou, A. H. A method to recapitulate early embryonic spatial patterning in human embryonic stem cells. Nat. Methods 11, 847–854 (2014).

3. Morgani, S. M., Metzger, J. J., Nichols, J., Siggia, E. D. & Hadjantonakis, A.-K. Micropattern differentiation of mouse pluripotent stem cells recapitulates embryo regionalized cell fate patterning. Elife 7, 1040 (2018).

4. Berge ten, D. et al. Wnt signaling mediates self-organization and axis formation in embryoid bodies. Cell Stem Cell 3, 508–518 (2008).

5. van den Brink, S. C. et al. Symmetry breaking, germ layer specification and axial organisation in aggregates of mouse embryonic stem cells. Development 141, 4231–4242 (2014).

6. Beccari, L. et al. Multi-axial self-organization properties of mouse embryonic stem cells into gastruloids. Nature 562, 272–276 (2018).

7. Rossi, G. et al. Capturing Cardiogenesis in Gastruloids. Cell Stem Cell (2020). doi:10.1016/j.stem.2020.10.013

8. Veenvliet, J. V. et al. Mouse embryonic stem cells self-organize into trunk-like structures with neural tube and somites. Science 370, eaba4937–9 (2020).

9. van den Brink, S. C. et al. Single-cell and spatial transcriptomics reveal somitogenesis in gastruloids. Nature 582, 405–409 (2020).

10. Rivron, N. C. et al. Blastocyst-like structures generated solely from stem cells. Nature 557, 106–111 (2018).

11. Harrison, S. E., Sozen, B., Christodoulou, N., Kyprianou, C. & Zernicka-Goetz, M. Assembly of embryonic and extraembryonic stem cells to mimic embryogenesis in vitro. Science 356, eaal1810 (2017).

12. Sozen, B. et al. Self-assembly of embryonic and two extra-embryonic stem cell types into gastrulating embryo-like structures. Nature Cell Biology 20, 979–989 (2018).

13. Brandenberg, N. et al. High-throughput automated organoid culture via stem-cell aggregation in microcavity arrays. Nat Biomed Eng 345, 1247125–12 (2020).

14. Eiraku, M. et al. Self-organizing optic-cup morphogenesis in three-dimensional culture. Nature 472, 51–56 (2011).

15. Sheng, G. Epiblast morphogenesis before gastrulation. Developmental Biology 401, 17–24 (2015).

16. Bedzhov, I. & Zernicka-Goetz, M. Self-Organizing Properties of Mouse Pluripotent Cells Initiate Morphogenesis upon Implantation. Cell 156, 1032–1044 (2014).

17. Meng, Y. et al. Pten facilitates epiblast epithelial polarization and proamniotic lumen formation in early mouse embryos. Dev. Dyn. 246, 517–530 (2017).

18. Turner, D. A. et al. Anteroposterior polarity and elongation in the absence of extra-embryonic tissues and of spatially localised signalling in gastruloids: mammalian embryonic organoids. Development 144, 3894–3906 (2017).

19. Donnison, M. et al. Loss of the extraembryonic ectoderm in Elf5 mutants leads to defects in embryonic patterning. Development 132, 2299–2308 (2005).

20. Latos, P. A. et al. Elf5-centered transcription factor hub controls trophoblast stem cell self-renewal and differentiation through stoichiometry-sensitive shifts in target gene networks. Genes & Development 29, 2435–2448 (2015).

21. Latos, P. A. & Hemberger, M. From the stem of the placental tree: trophoblast stem cells and their progeny. Development 143, 3650–3660 (2016).

22. Guzman-Ayala, M., Ben-Haim, N., Beck, S. & Constam, D. B. Nodal protein processing and fibroblast growth factor 4 synergize to maintain a trophoblast stem cell microenvironment. Proceedings of the National Academy of Sciences 101, 15656–15660 (2004).

23. Rivera-Pérez, J. A. & Hadjantonakis, A.-K. The Dynamics of Morphogenesis in the Early Mouse Embryo. Cold Spring Harbor Perspectives in Biology 7, a015867 (2014).

24. Kubaczka, C. et al. Derivation and maintenance of murine trophoblast stem cells under defined conditions. Stem Cell Reports 2, 232–242 (2014).

25. Rodriguez, T. A., Srinivas, S., Clements, M. P., Smith, J. C. & Beddington, R. S. P. Induction and migration of the anterior visceral endoderm is regulated by the extra-embryonic ectoderm. Development 132, 2513–2520 (2005).

26. Winnier, G., Blessing, M., Labosky, P. A. & Hogan, B. L. Bone morphogenetic protein-4 is required for mesoderm formation and patterning in the mouse. Genes & Development 9, 2105–2116 (1995).

27. Hoffman, J. A., Wu, C.-I. & Merrill, B. J. Tcf7l1 prepares epiblast cells in the gastrulating mouse embryo for lineage specification. Development 140, 1665–1675 (2013).

28. Wolfe, A. D., Rodriguez, A. M. & Downs, K. M. STELLA collaborates in distinct mesendodermal cell subpopulations at the fetal-placental interface in the mouse gastrula. Developmental Biology 425, 44–57 (2017).

29. Rossant, J. & Tam, P. P. L. Blastocyst lineage formation, early embryonic asymmetries and axis patterning in the mouse. Development 136, 701–713 (2009).

30. Huelsken, J. et al. Requirement for beta-catenin in anterior-posterior axis formation in mice. The Journal of Cell Biology 148, 567–578 (2000).

31. Ben-Haim, N. et al. The nodal precursor acting via activin receptors induces mesoderm by maintaining a source of its convertases and BMP4. Developmental Cell 11, 313–323 (2006).

32. Ferrer-Vaquer, A. et al. A sensitive and bright single-cell resolution live imaging reporter of Wnt/ß-catenin signaling in the mouse. BMC Dev. Biol. 10, 121–18 (2010).

33. Faunes, F. et al. A membrane-associated β-catenin/Oct4 complex correlates with ground-state pluripotency in mouse embryonic stem cells. Development 140, 1171–1183 (2013).

34. Serup, P. et al. Partial promoter substitutions generating transcriptional sentinels of diverse signaling pathways in embryonic stem cells and mice. Dis Model Mech 5, 956–966 (2012).

35. Cajal, M. et al. Clonal and molecular analysis of the prospective anterior neural boundary in the mouse embryo. Development 139, 423–436 (2012).

36. Li, L. et al. Location of transient ectodermal progenitor potential in mouse development. Development 140, 4533–4543 (2013).

37. Lewis, S. L. et al. Dkk1 and Wnt3 interact to control head morphogenesis in the mouse. Development 135, 1791–1801 (2008).

38. Perea-Gomez, A. et al. Nodal antagonists in the anterior visceral endoderm prevent the formation of multiple primitive streaks. Developmental Cell 3, 745–756 (2002).

39. Girgin, M. U. & Lutolf, M. P. Gastruloids generated without exogenous Wnt activation develop anterior neural tissues. bioRxiv 3, 2020.10.10.334326 (2020).

40. Maretto, S. et al. Mapping Wnt/beta-catenin signaling during mouse development and in colorectal tumors. Proceedings of the National Academy of Sciences 100, 3299–3304 (2003).

41. Fausett, S. R., Brunet, L. J. & Klingensmith, J. BMP antagonism by Noggin is required in presumptive notochord cells for mammalian foregut morphogenesis. Developmental Biology 391, 111–124 (2014).

42. Dessaud, E., McMahon, A. P. & Briscoe, J. Pattern formation in the vertebrate neural tube: a sonic hedgehog morphogen-regulated transcriptional network. Development 135, 2489–2503 (2008).

43. Beck, S. et al. Extraembryonic proteases regulate Nodal signalling during gastrulation. Nature Cell Biology 4, 981–985 (2002).

44. Ohinata, Y. & Tsukiyama, T. Establishment of trophoblast stem cells under defined culture conditions in mice. PLoS ONE 9, e107308 (2014).

45. Parker, H. J. & Krumlauf, R. Segmental arithmetic: summing up the Hox gene regulatory network for hindbrain development in chordates. Wiley Interdiscip Rev Dev Biol 6, e286 (2017).

46. Delile, J. et al. Single cell transcriptomics reveals spatial and temporal dynamics of gene expression in the developing mouse spinal cord. Development 146, dev173807 (2019).

47. Zhang, S. et al. Implantation initiation of self-assembled embryo-like structures generated using three types of mouse blastocyst-derived stem cells. Nature Communications 10, 1–17 (2019).

48. Etoc, F. et al. A Balance between Secreted Inhibitors and Edge Sensing Controls Gastruloid Self-Organization. Developmental Cell 39, 302–315 (2016).

49. Zhang, Z., Zwick, S., Loew, E., Grimley, J. S. & Ramanathan, S. Mouse embryo geometry drives formation of robust signaling gradients through receptor localization. Nature Communications 10, 4516–14 (2019).

50. Lancaster, M. A. et al. Cerebral organoids model human brain development and microcephaly. Nature 501, 373–379 (2013).

51. Qian, X. et al. Brain-Region-Specific Organoids Using Mini-bioreactors for Modeling ZIKV Exposure. Cell 165, 1238–1254 (2016).

52. Noémie M. L. P. Bérenger-Currias et al. Early neurulation recapitulated in assemblies of embryonic and extraembryonic cells. bioRxiv 11, 2020.02.13.947655 (2020).

53. Lewis, S. L. et al. Dkk1 and Wnt3 interact to control head morphogenesis in the mouse. Development 135, 1791–1801 (2008).

54. Stuckey, D. W., Di-Gregorio, A., Clements, M. & Rodriguez, T. A. Correct patterning of the primitive streak requires the anterior visceral endoderm. PLoS ONE 6, e17620 (2011).

55. Deluz, C. et al. A role for mitotic bookmarking of SOX2 in pluripotency and differentiation. Genes & Development 30, 2538–2550 (2016).

56. Brandenberg, N. et al. High-throughput automated organoid culture via stem-cell aggregation in microcavity arrays. Nat Biomed Eng 345, 1–12 (2020).

57. Martin, M. Cutadapt removes adapter sequences from high-throughput sequencing reads. EMBnet.journal 17, 10–12 (2011).

58. Stuart, T. et al. Comprehensive Integration of Single-Cell Data. Cell 177, 1888–1902.e21 (2019).

59. Pijuan-Sala, B. et al. A single-cell molecular map of mouse gastrulation and early organogenesis. Nature 566, 490–495 (2019).

60. Kiselev, V. Y., Yiu, A. & Hemberg, M. scmap: projection of single-cell RNA-seq data across data sets. Nat. Methods 15, 359–362 (2018).

61. McGinnis, C. S. et al. MULTI-seq: sample multiplexing for single-cell RNA sequencing using lipid-tagged indices. Nat. Methods 16, 619–626 (2019).

62. Korsunsky, I. et al. Fast, sensitive and accurate integration of single-cell data with Harmony. Nat. Methods 16, 1289–1296 (2019).

63. Becht, E. et al. Dimensionality reduction for visualizing single-cell data using UMAP. Nature Biotechnology 37, 38–44 (2018).

64. La Manno, G. et al. RNA velocity of single cells. Nature 560, 494–498 (2018).

65. Bergen, V., Lange, M., Peidli, S., Wolf, F. A. & Theis, F. J. Generalizing RNA velocity to transient cell states through dynamical modeling. Nature Biotechnology 38, 1408–1414 (2020).

66. Wolf, F. A., Angerer, P. & Theis, F. J. SCANPY: large-scale single-cell gene expression data analysis. Genome Biol 19, 15–5(2018).

