## Supplementary Data for "Bioengineered embryoids mimic post-implantation development *in vitro*"

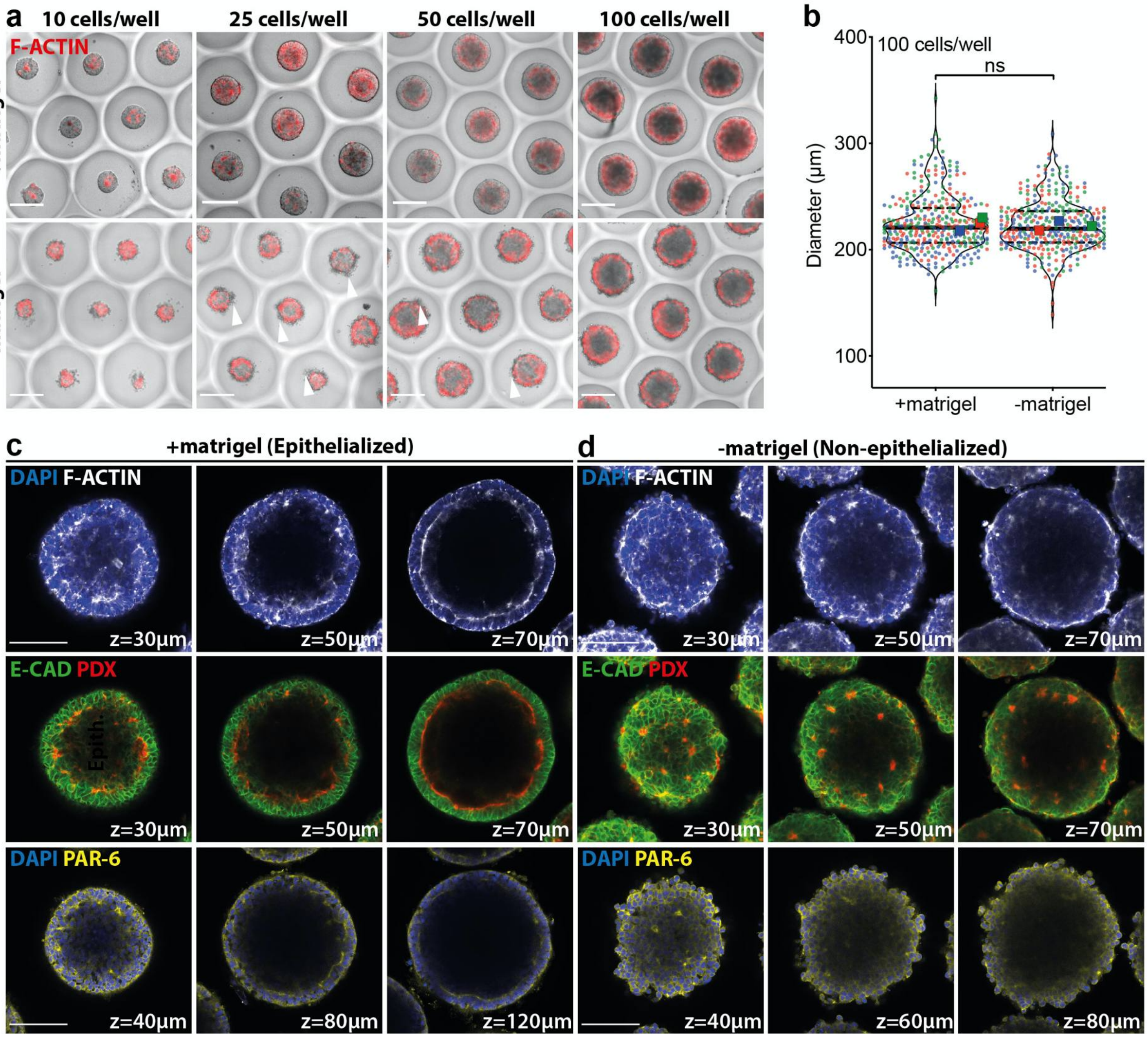


**Supplementary Figure 1: Effects of starting cell number and addition of Matrigel on epithelialization of EPI aggregates. a)** Representative confocal images at 72 h showing the effects of starting cell number and addition of Matrigel on *F-actin* expression (phalloidin staining). Note the shed cells around the aggregates (white arrows) in the absence of Matrigel. **b)** Comparing minimum ferret diameters of EPI aggregates at 72 h formed with or without Matrigel from 100 cells/well. For +matrigel and -matrigel conditions, total number of aggregates analyzed were 361 and 348, respectively. Data is collected from three biologically independent experiments. Large symbols indicate mean values of each replicate. Black lines indicate median and quartiles. **c,d)** Confocal images of showing multiple z-planes of EPI aggregates formed from 100 cells/well with **(c)** or without **(d)** Matrigel fixed at 72 h and stained for *F-actin* (phalloidin), *E-cadherin*, *Podocalxyin* and *Par6*. Nuclei were stained with DAPI. For statistical analysis, two-tailed unpaired Student's t-test **(b)** was performed. Following P-value style was used: ns=not significant, P****<0.0001, P***<0.0002, P**<0.0021, P*<0.0332. Scale bars: 100µm.

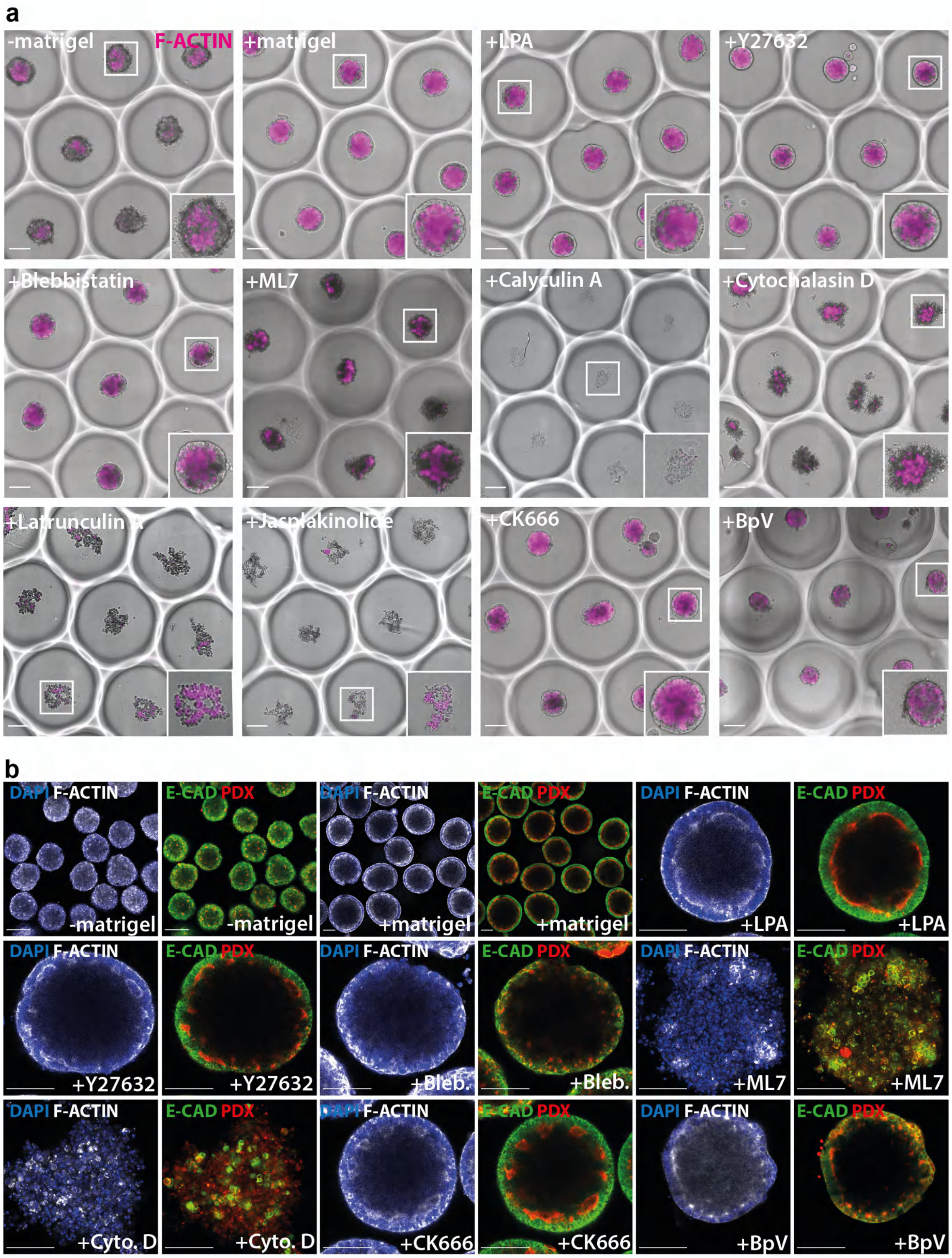


**Supplementary Figure 2: Effect of cytoskeleton inhibitors on epithelialization of EPI aggregates. a)** Representative images showing *F-actin* (phalloidin) expression in EPI aggregates on microwells, cultured with Matrigel and indicated inhibitors for 72 h. **b)** Representative confocal images of EPI aggregates cultured with Matrigel and indicated inhibitors for 72 h, showing *E-cadherin, Podocalyxin* and *F-actin* (phalloidin) expression. Scale bars: 100µm.

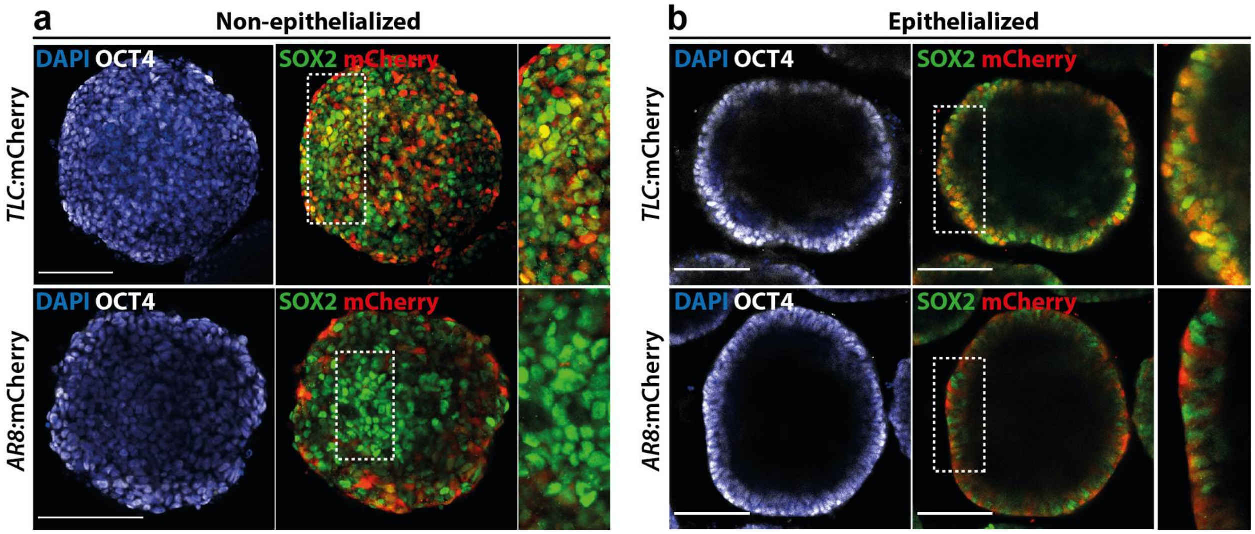


**Supplementary Figure 3: WNT and TGF-βsignaling in EPI aggregates. a,b)** Representative confocal images showing *Oct4, Sox2* expression and WNT *(TLC:mCherry*, top) or TGF-β (*AR8:mCherry*, bottom) reporter activity in non-epithelialized **(a)** or epithelialized **(b)** EPI aggregates at 72 h. Scale bars: 100μm.

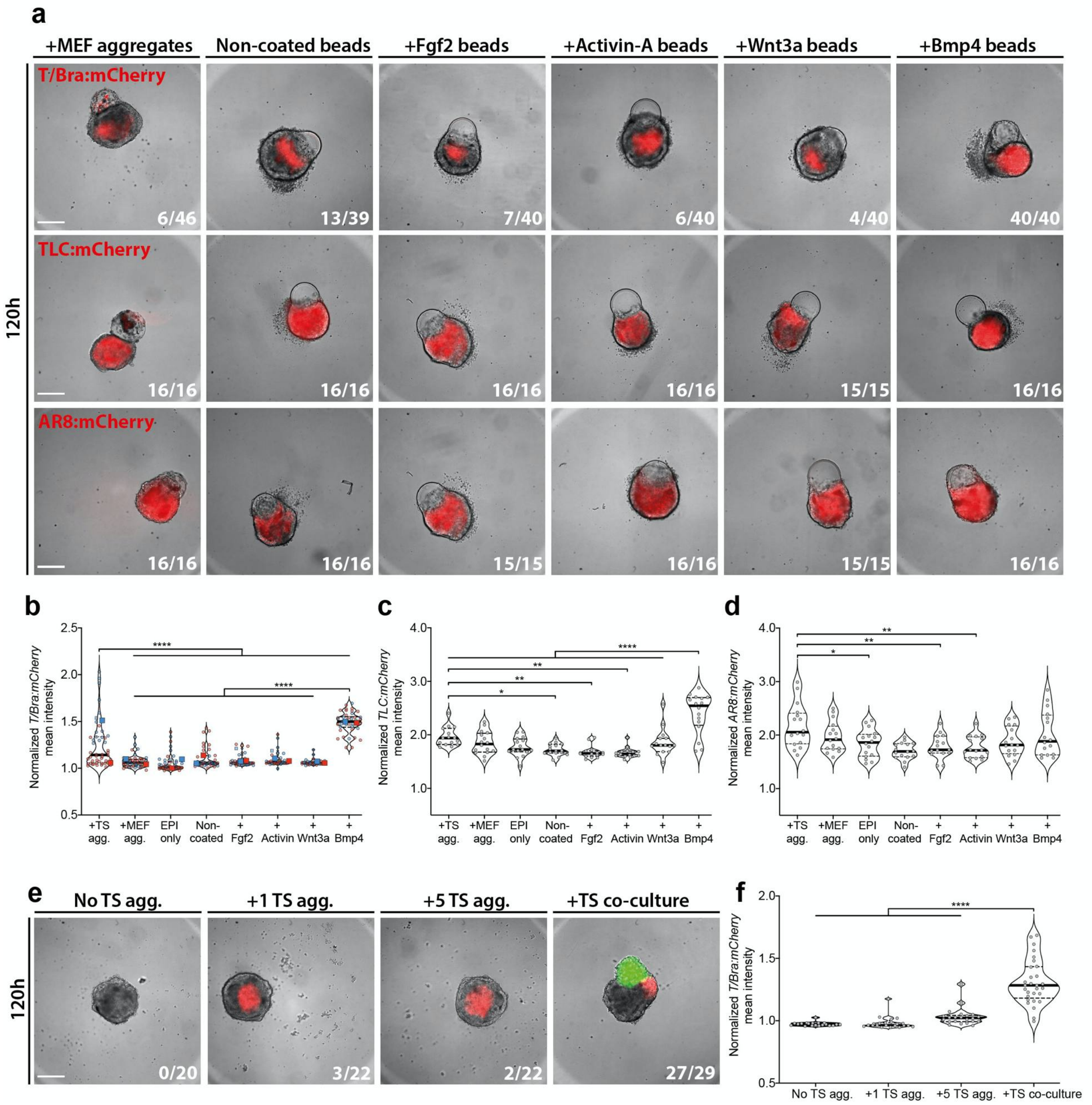


**Supplementary Figure 4: Dependence of *T/Bra* expression in epithelialized EPI aggregates to TS aggregate derived signaling a)** Representative images showing *T/Bra*:mCherry (top), *TLC*:mCherry (middle) and *AR8:*mCherry (bottom) reporter activities at 120h in epithelialized EPI aggregates co-cultured with mouse embryonic fibroblast (MEF) aggregates or beads coated with indicated proteins. After transfer, the medium was not changed until 120h in order to concentrate the factors. **b-d)** Quantification of background normalized mean intensities of *T/Bra*:mCherry **(b)**, *TLC*:mCherry **(c)** and *AR8:*mCherry **(d)** at 120h in epithelialized EPI aggregates co-cultured in indicated conditions. Data was collected from two independent experiments **(b)** or from single experiments **(c,d)**. **e)** Representative images showing *T/Bra* expression in epithelialized EPI aggregates co-cultured with indicated number of TS aggregates on transwells. **f)** Quantification of background normalized mean intensity of *T/Bra*:mCherry in transwell co-culture at 120h. Data was collected from single experiment. For all conditions in **(a)** and **(c)**, number of *T/Bra*:mCherry -positive embryoids over total number of embryoids analyzed are indicated at bottom right. For **(b)** and **(d)**, large symbols indicate mean values of each replicate. Black lines indicate median and quartiles. For all statistical analysis, one-way ANOVA followed by Tukey multiple comparison test was performed. Following P-value style was used: P****<0.0001, P***<0.0002, P**<0.0021, P*<0.0332. Scale bars: 200μm.

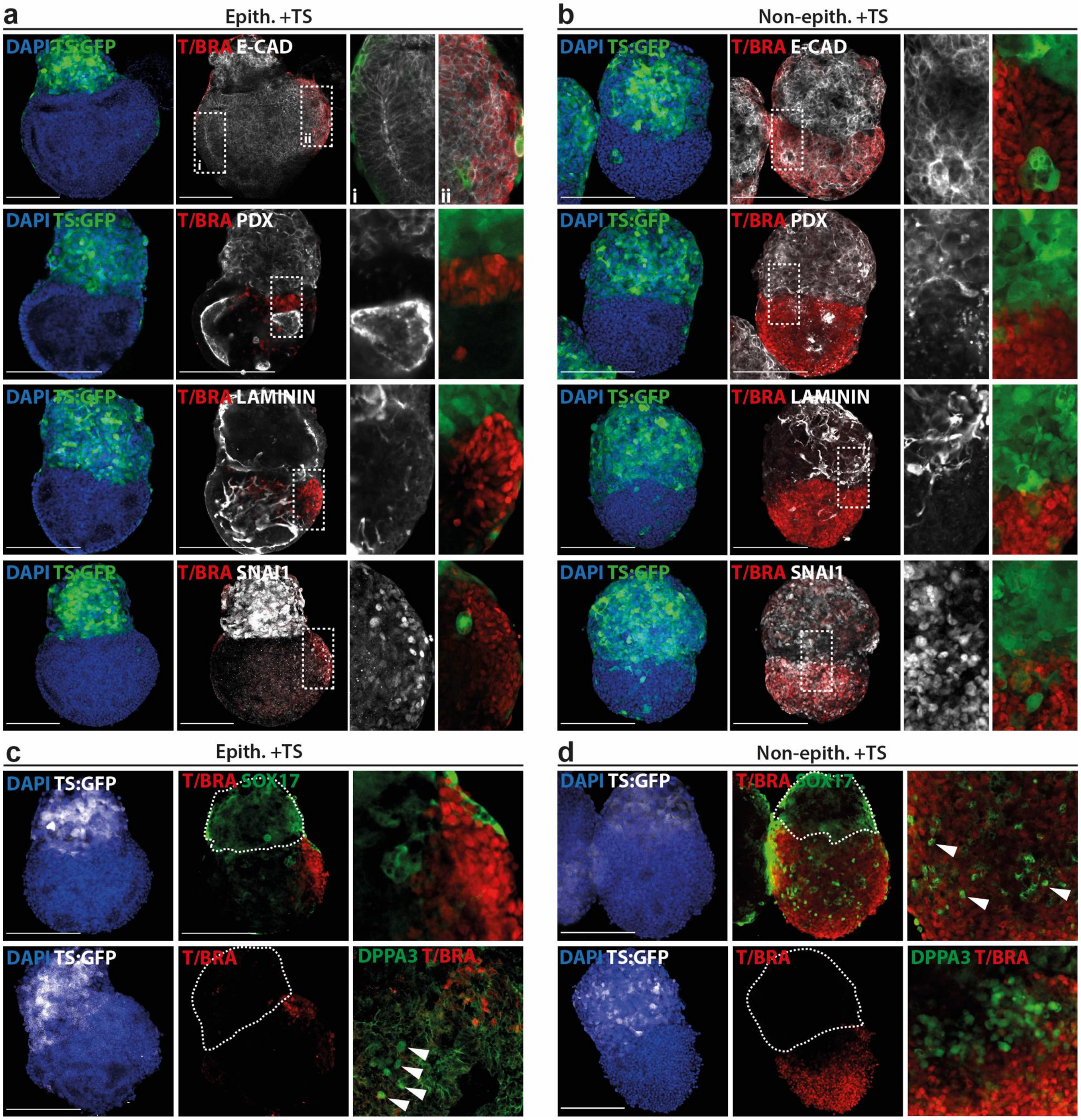


**Supplementary Figure 5: Characterization of epiblast and gastrulation markers in epithelialized and non-epithelialized embryoids at 120h. a-b)** Representative confocal images showing *E-cadherin, Podocalyxin, Laminin* and *Snai1* expression in epithelialized **(a)** and non-epithelialized **(b)** embryoids. **c-d)** Representative confocal images showing *Sox17, Stella* and *T/Bra* expression in epithelialized **(c)** and non-epithelialized **(d)** embryoids. Nuclei were stained with DAPI. GFP-labeled TS cells were depicted in green **(a,b)** or in white (**c,d)**.

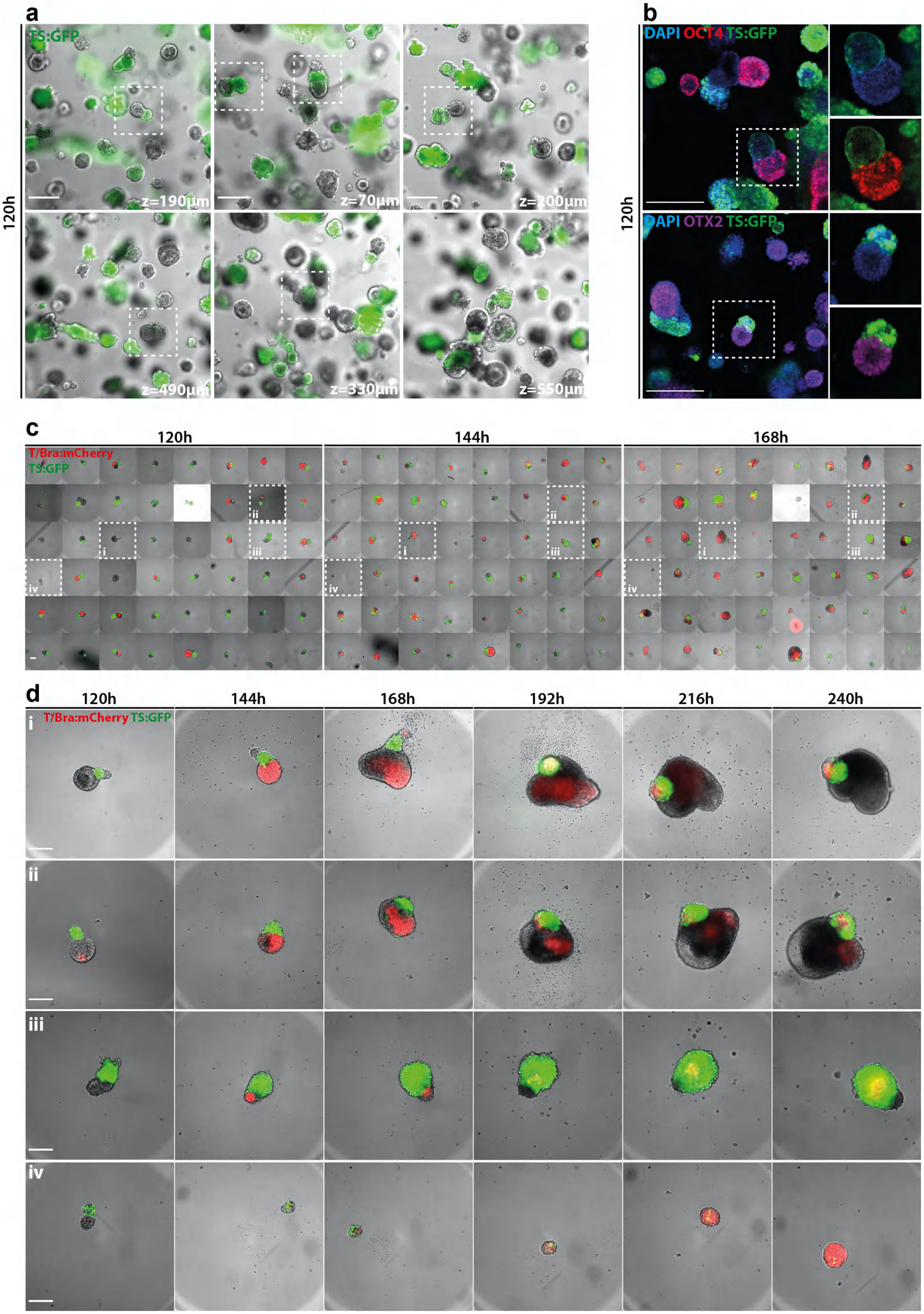

a
TS:GFP
120h
z=190µm
z=70µm
z=200µm
z=490µm
z=330µm
z=550µm
b
DAPI OCT4 TS:GFP
DAPI OTX2 TS:GFP
120h
c
120h
144h
168h
T/Bra:mCherry
TS:GFP
d
120h
144h
168h
192h
216h
240h
T/Bra:mCherry TS:GFP
i
ii
iii
iv

**Supplementary Figure 6: Long term culture of ETS embryos a)** Representative images showing ETS embryos in culture at 120h. **b)** Representative confocal images showing *Oct4* and *Otx2* immunostainings at 120h. **c**) Montage of ETS embryos transferred to 96 well plate on 120h and cultured until 168h. **d)** Timepoint images showing extended culture until 240h and representative phenotypes.

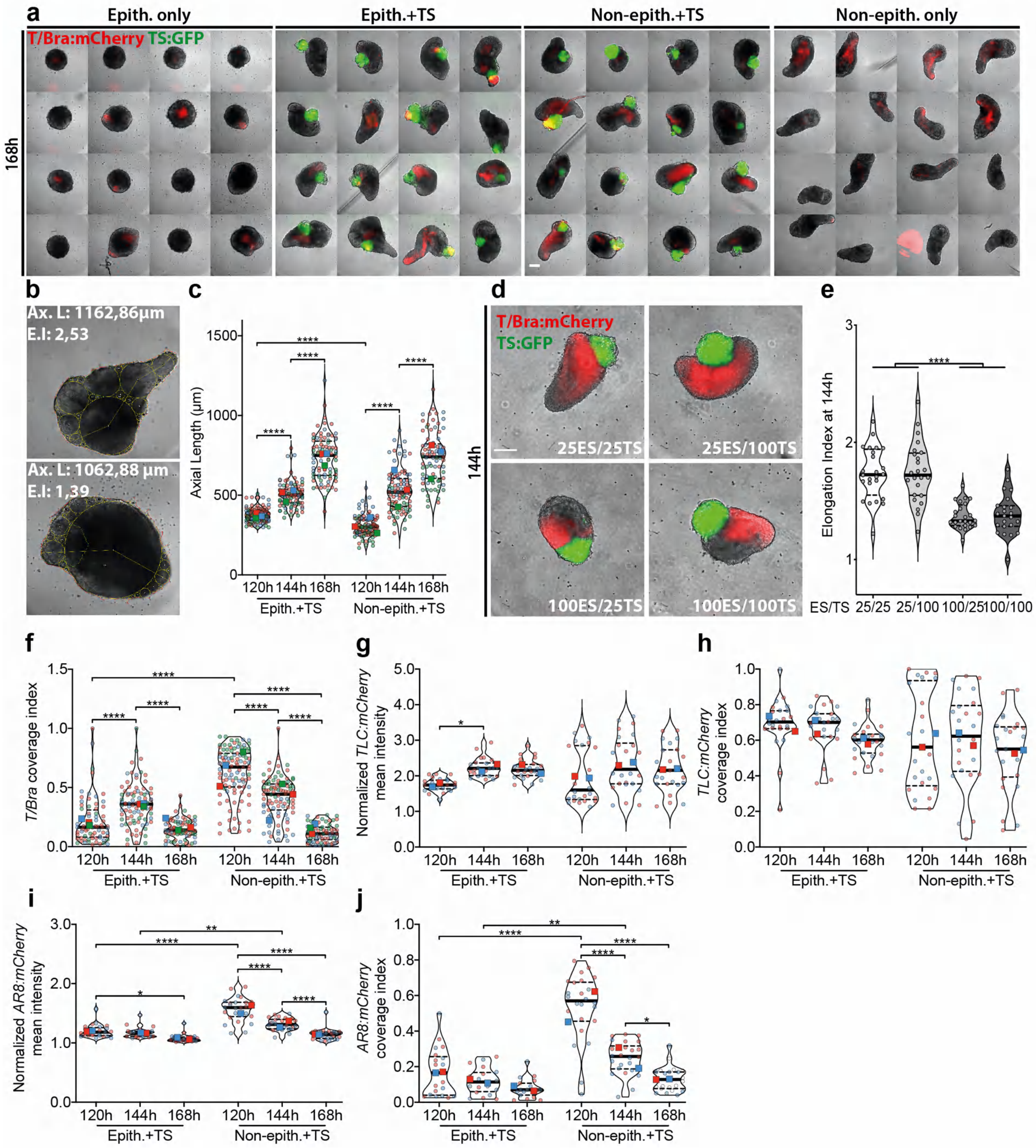


**Supplementary Figure 7: Axial morphogenesis dynamics EpiTS embryoids a)** Montage view of a single experiment showing reproducibility in morphology within each condition. **b)** Representative images showing axial length and elongation index calculation. **c)** Quantification of axial length of epithelialized and non-epithelialized embryoids between 120h to 168h. **d-e)** Representative images **(d)** and quantification of elongation index **(e)** of epithelialized embryoids formed from 25/25, 25/100, 100/25 and 100/100 conditions at 144h. Total number of embryoids analyzed were 24, 24, 24 and 23, respectively. For elongation index quantification, TS subtraction was performed. **f-j)** Quantification of coverage index of *T/Bra:*mCherry **(f),** *TLC:*mCherry **(h)**, *AR8:*mCherry **(j)** and background normalized *TLC:*mCherry **(g)**, *AR8:*mCherry **(i)** mean intensity in epithelialized or non-epithelialized embryoids between 120h to 168h. For all conditions in **(c,f-j)**, total number of embryoids analyzed are indicated at bottom right of **(Figure 4a-b)**. Large symbols indicate mean values of each replicate. Black lines indicate median and quartiles. For all statistical analysis, one-way ANOVA followed by Tukey multiple comparison test was performed. Following P-value style was used: P****<0.0001, P***<0.0002, P**<0.0021, P*<0.0332. Scale bars: 200µm.

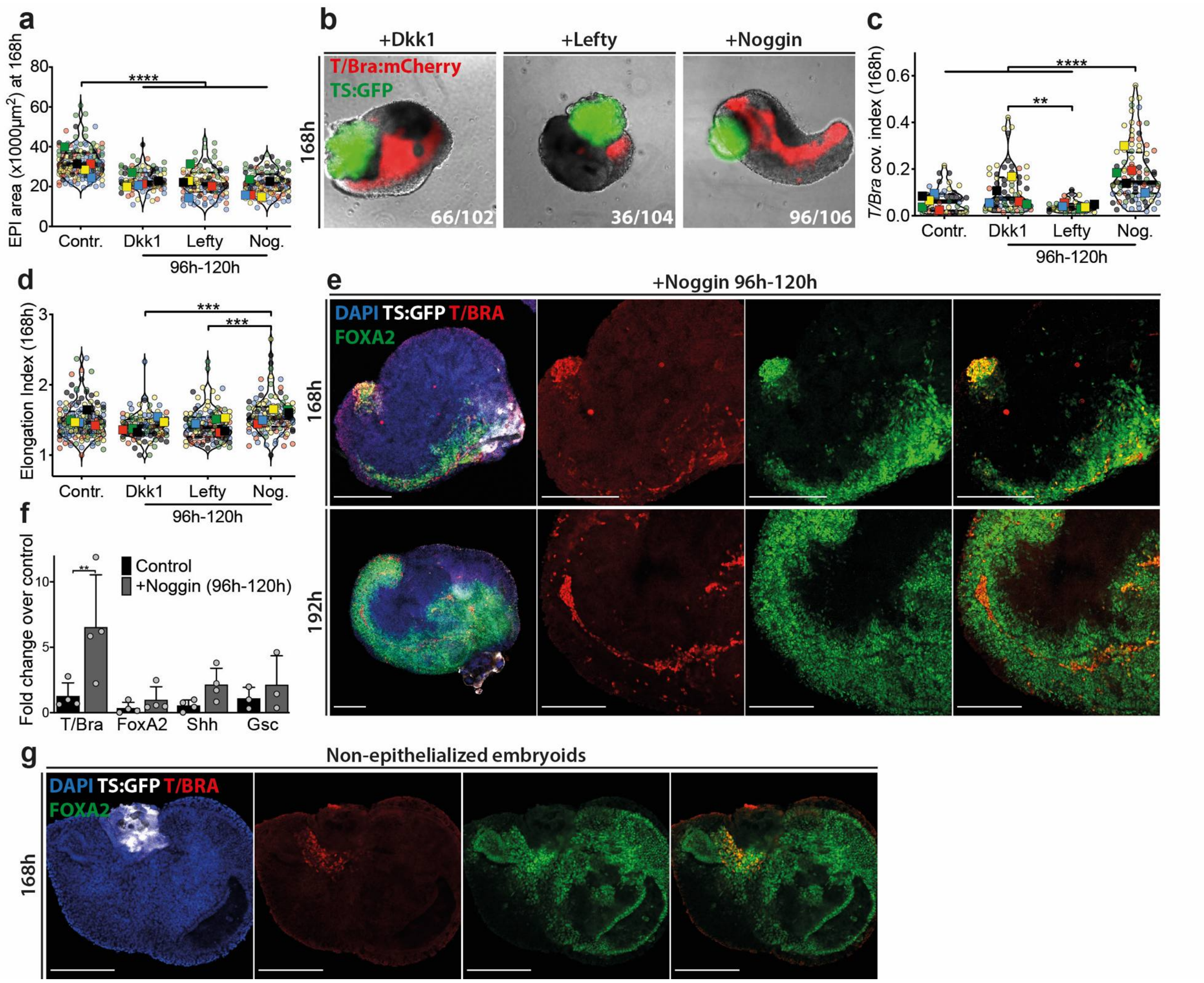


**Supplementary Figure 8: Roles of WNT, TGF-β and BMP pathways in axial morphogenesis a)** Quantification of EPI compartment area at 168h in epithelialized embryoids treated with indicated inhibitors between 96h-120h. **b)** Representative images at 168h showing *T/Bra:*mCherry expression in epithelialized embryoids treated with indicated inhibitors between 96h to 120h. **b-c)** Quantification of coverage index of *T/Bra:*mCherry expression **(b)** and elongation index **(c)** at 168h in epithelialized embryoids treated with indicated inhibitors. For elongation index quantification, TS subtraction was not performed. **e)** Representative confocal images of Noggin-treated epithelialized embryoid showing *T/Bra* and *Foxa2* immunostainings at 168h and 192h. **f)** RT-PCR analysis showing expression levels of *T/Bra, Foxa2, Shh* and *Gsc* in Noggin-treated epithelialized embryoids compared to untreated embryoids. Data is shown as mean. **g)** Representative confocal images of non-epithelialized embryoids showing *T/Bra* and *Foxa2* immunostainings at 168h. Nuclei were stained with DAPI. For all conditions in **(a,c,d)** total number of embryoids analyzed are indicated at bottom right of **(b)**. Large symbols indicate mean values of each replicate. Black lines indicate median and quartiles. For all statistical analysis, one-way ANOVA followed by Tukey multiple comparison test was performed. Following P-value style was used: P****<0.0001, P***<0.0002, P**<0.0021, P*<0.0332. Scale bars: 200µm.

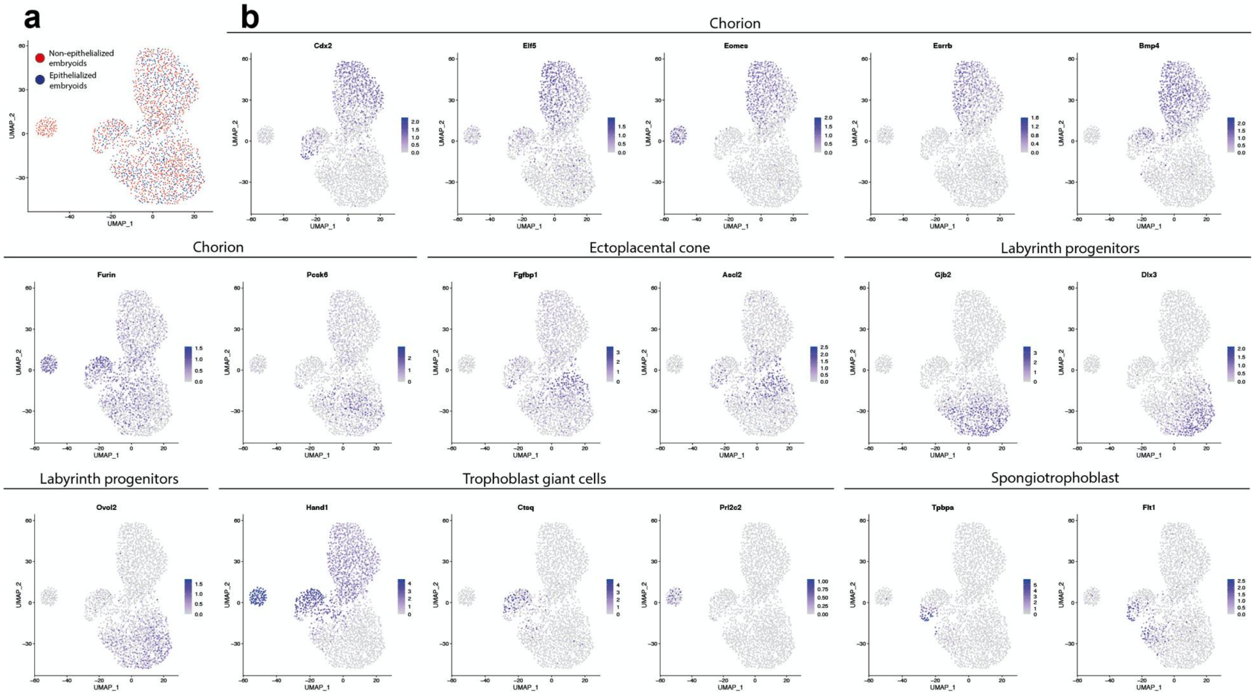


**Supplementary Figure 9: Expression of key extraembryonic tissue markers in EpiTS embryoids. a)** Demonstration of sample origin of cell types in extraembryonic cluster. **b)** Key markers for each extraembryonic cell type observed in EpiTS embryoids.

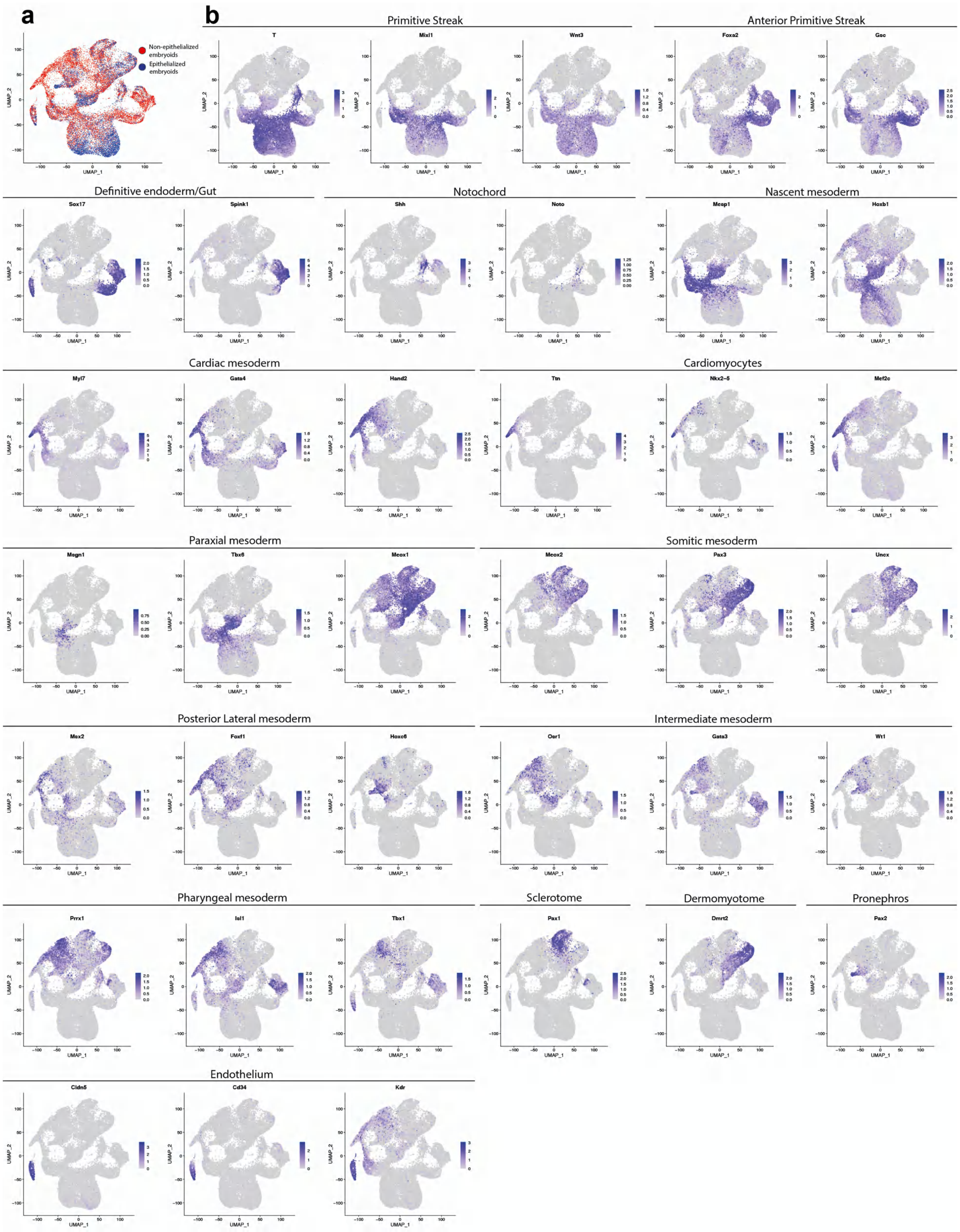


**Supplementary Figure 10: Expression of key mesendoderm markers in EpiTS embryoids.** **a)** Demonstration of sample origin of cell types in mesendoderm cluster. **b)** Key markers for each mesendoderm cell type observed in EpiTS embryoids.

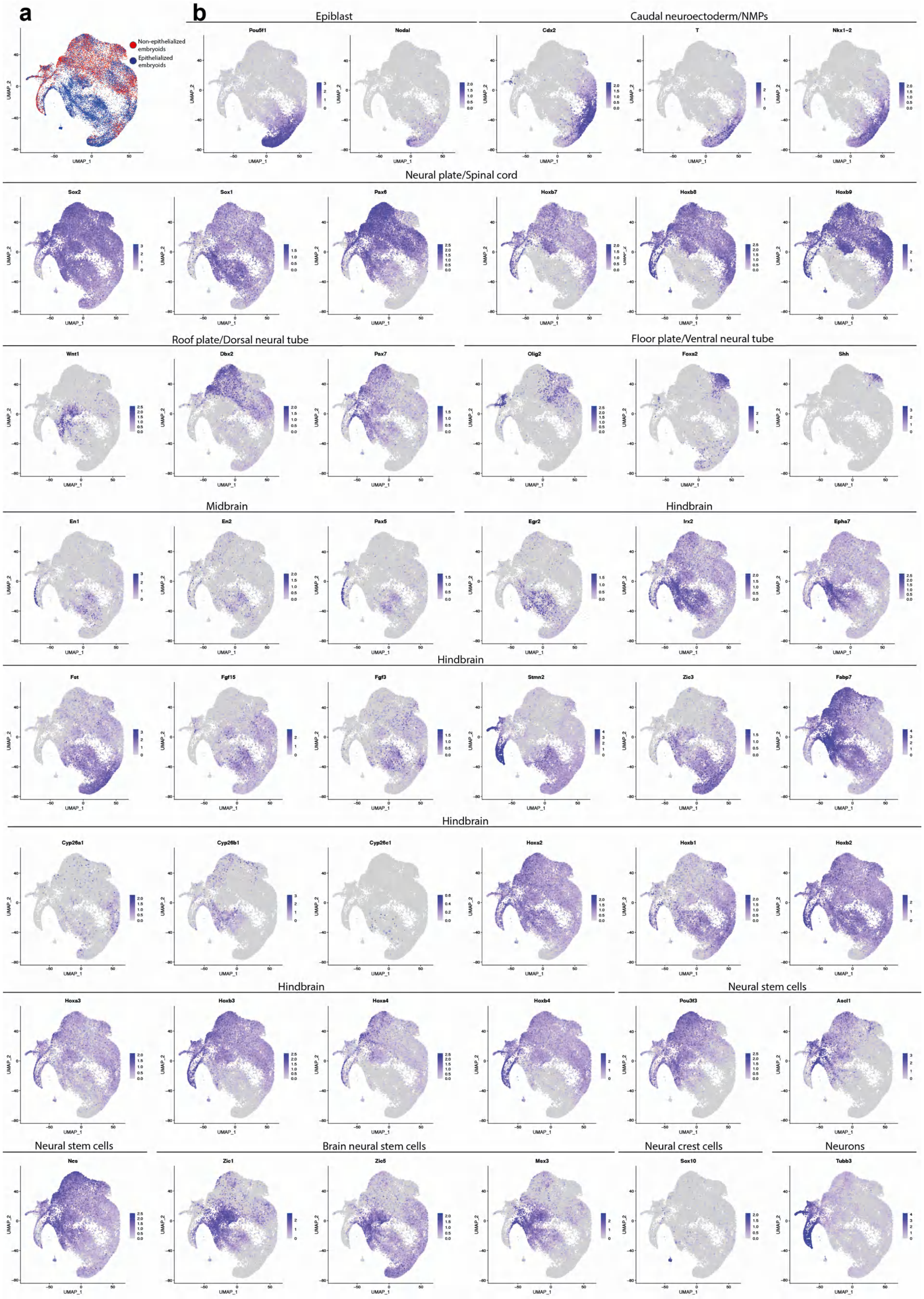
a
Non-epithelialized embryoids
Epithelialized embryoids
b
Epiblast
Caudal neuroectoderm/NMPs
Pou5f1
Nodal
Cdx2
T
Nkx1-2
Neural plate/Spinal cord
Sox2
Sox1
Pax6
Hoxb7
Hoxb8
Hoxb9
Roof plate/Dorsal neural tube
Floor plate/Ventral neural tube
Wnt1
Dbx2
Pax7
Olig2
Foxa2
Shh
Midbrain
Hindbrain
En1
En2
Pax5
Egr2
Irx2
Epha7
Hindbrain
Fst
Fgf15
Fgf3
Stmn2
Zic3
Fabp7
Hindbrain
Cyp26a1
Cyp26b1
Cyp26c1
Hoxa2
Hoxb1
Hoxb2
Hindbrain
Neural stem cells
Hoxa3
Hoxb3
Hoxa4
Hoxb4
Pou3f3
Ascl1
Neural stem cells
Brain neural stem cells
Neural crest cells
Neurons
Nes
Zic1
Zic5
Msx3
Sox10
Tubb3
UMAP_1
UMAP_2

**Supplementary Figure 11: Expression of key epiblast and ectoderm markers in EpiTS embryoids. a)** Demonstration of sample origin of cell types in ectoderm cluster. **b)** Key markers for each ectoderm cell type observed in EpiTS embryoids.

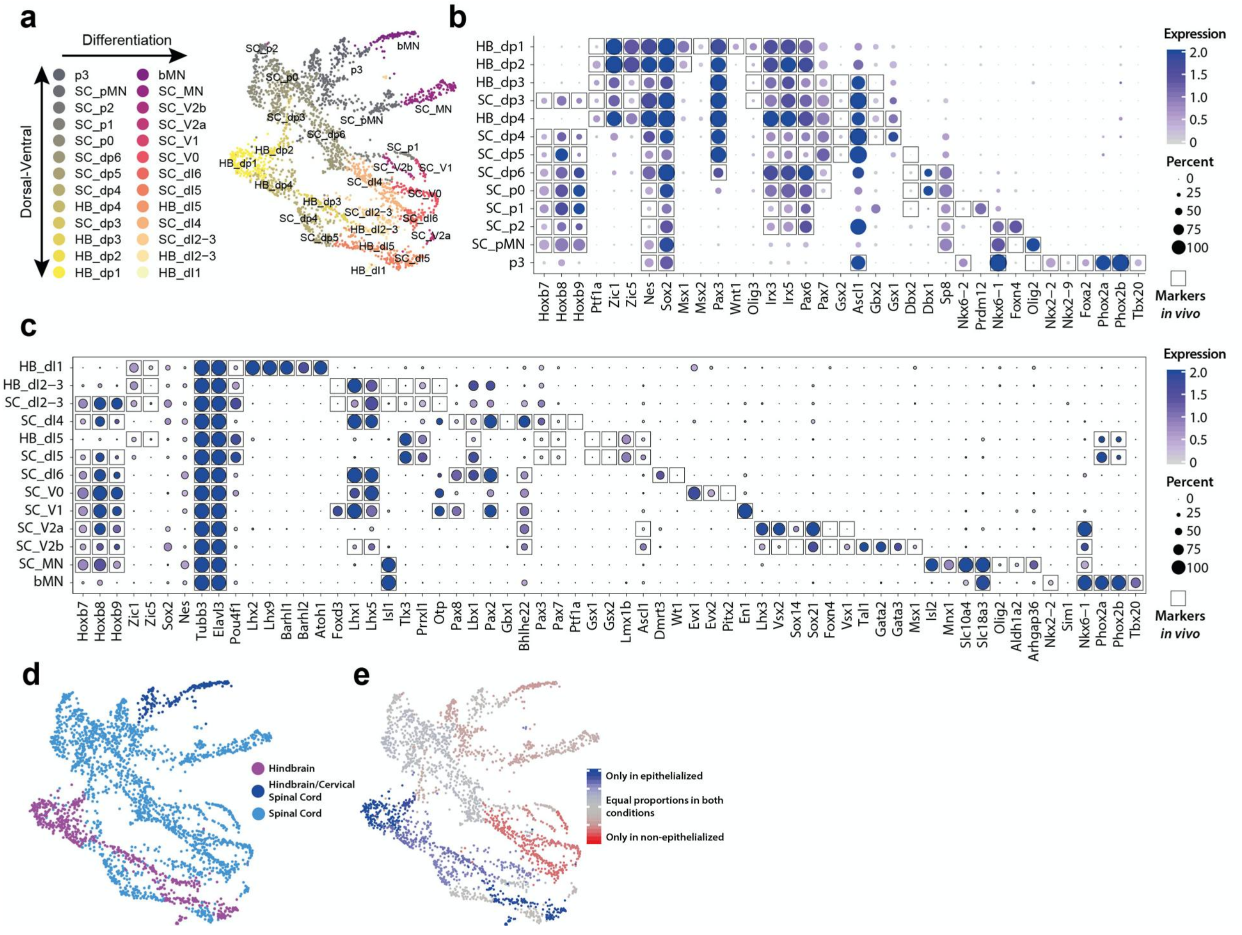


**Supplementary Figure 12: Diversification of neural progenitors and neurons in EpiTS embryoids. a)** Demonstration of cell types in neurons cluster mapping to dorsal-ventral axis **b-c)** Key markers for neural progenitor **(b)** and neurons **(c)** with reference to *in vivo* atlas. **d-e)** Demonstration of tissue origin **(d)** or sample origin **(e)** of cell types in neuron cluster.

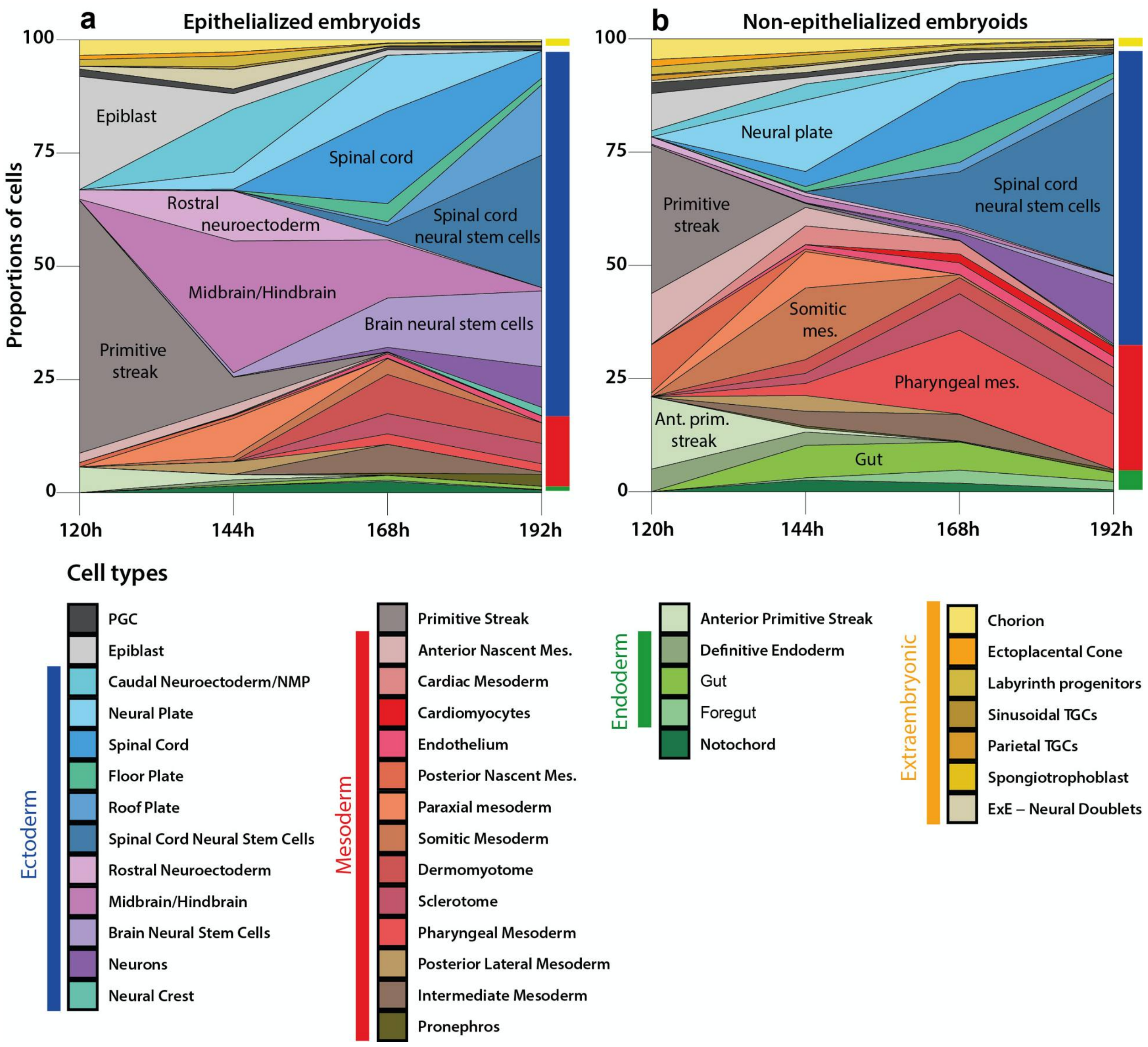


**Supplementary Figure 13: Cell type proportion in epithelialized and non-epithelialized embryoids. a,b)** Percentage of cells that constitute indicated cell populations between 120h to 192h in epithelialized **(a)** or non-epithelialized **(b)** embryoids.

### Supplementary Movies

**Supplementary Movie 1:** Timelapse imaging showing *T/Bra* expression dynamics until 144h in *EpiTS embryoids* formed from 25ESC/25TSC condition. Images were acquired every 2 hours. *T/Bra* expression was depicted in red, TS:GFP cells were shown in green. Scale bar: 200µm.

**Supplementary Movie 2:** Timelapse imaging showing *T/Bra* expression dynamics until 144h in *EpiTS embryoids* formed from 25ESC/100TSC condition. Images were acquired every 2 hours. *T/Bra* expression was depicted in red, TS:GFP cells were shown in green. Scale bar: 200µm.

**Supplementary Movie 3:** Timelapse imaging showing *T/Bra* expression dynamics until 144h in *EpiTS embryoids* formed from 100ESC/25TSC condition. Images were acquired every 2 hours. *T/Bra* expression was depicted in red, TS:GFP cells were shown in green. Scale bar: 200µm.

**Supplementary Movie 4:** Timelapse imaging showing *T/Bra* expression dynamics until 144h in *EpiTS embryoids* formed from 100ESC/100TSC condition. Images were acquired every 2 hours. *T/Bra* expression was depicted in red, TS:GFP cells were shown in green. Scale bar: 200µm.

**Supplementary Movie 5:** Timelapse imaging showing *T/Bra* expression dynamics until 160h in epithelialized *EpiTS embryoids* formed from 100ESC/100TSC condition. Images were acquired every 2 hours. *T/Bra* expression was depicted in red, TS:GFP cells were shown in green. Scale bar: 200µm.

**Supplementary Movie 6:** Timelapse imaging showing *T/Bra* expression dynamics until 160h in non-epithelialized *EpiTS embryoids* formed from 100ESC/100TSC condition. Images were acquired every 2 hours. *T/Bra* expression was depicted in red, TS:GFP cells were shown in green. Scale bar: 200µm.

**Supplementary Movie 7:** Timelapse imaging showing *T/Bra* expression dynamics until 168h in epithelialied *EpiTS embryoids* formed from from 100ESC/100TSC condition. Images were acquired every 2 hours. *T/Bra* expression was depicted in red, TS:GFP cells were shown in green. Scale bar: 200µm.

**Supplementary Movie 8:** Timelapse imaging showing *T/Bra* expression dynamics until 168h in epithelialied *EpiTS embryoids* formed from from 100ESC/100TSC condition and treated with 200ng/ml Dkk1 between 96h-120h. Images were acquired every 2 hours. *T/Bra* expression was depicted in red, TS:GFP cells were shown in green. Scale bar: 200µm.

**Supplementary Movie 9:** Timelapse imaging showing *T/Bra* expression dynamics until 168h in epithelialied *EpiTS embryoids* formed from from 100ESC/100TSC condition and treated with 200ng/ml LeftyA between 96h-120h. Images were acquired every 2 hours. *T/Bra* expression was depicted in red, TS:GFP cells were shown in green. Scale bar: 200µm.

**Supplementary Movie 10:** Timelapse imaging showing *T/Bra* expression dynamics until 168h in epithelialied *EpiTS embryoids* formed from from 100ESC/100TSC condition and treated with 200ng/ml Noggin between 96h-120h. Images were acquired every 2 hours. *T/Bra* expression was depicted in red, TS:GFP cells were shown in green. Scale bar: 200µm.

**Table 1: List of primary antibodies used for immunostaining**

| Target | Species | Dilution | Catalogue Number | Supplier |
|---|---|---|---|---|
| anti-E-cadherin | Rabbit | 1:500 | #24E10 | Cell Signaling Technology |
| anti-Podocalyxin | Rat | 1:200 | #MAB1556 (192703) | R&D systems |
| anti-Par6 | Mouse | 1:100 | #sc-166405 (B-10) | Santa Cruz |
| anti- aPKC | Mouse | 1:100 | #sc-17781 (H-1) | Santa Cruz |
| anti-Sox1 | Goat | 1:50 | #af3369 | R&D Systems |
| anti-Sox2 | Rabbit | 1:400 | #ab97959 | Abcam |
| anti-Pax6 | Rabbit | 1:100 | #901301 (Poly19013) | BioLegend |
| anti-Otx2 | Goat | 1:25 | #af1979 | R&D Systems |
| anti-Tuj1 | RAbbit | 1:400 | #ab18207 | Abcam |
| anti-Brachyury | Goat | 1:300 | #sc-17745 (C-19) | Santa Cruz |
| anti-Brachyury | Rabbit | 1:100 | #ab209665 | Abcam |
| anti-Oct4 | Mouse | 1:200 | #sc-5270 (C-10) | Santa Cruz |
| anti-Nanog | Rat | 1:300 | #14-5761-80 | ThermoFisher |
| anti-Dppa3 | Mouse | 1:100 | #AF2566-SP | R&D systems |
| anti-Sox17 | Goat | 1:200 | #AF1924 | Abcam |
| anti-Foxa2 | Rabbit | 1:200 | #ab108422 | Abcam |
| anti-Cdx2 | Rabbit | 1:200 | #ab76541 | Abcam |
| anti-Eomes | Rabbit | 1:200 | #ab23345 | Abcam |
| anti-Tfap2c | Mouse | 1:200 | #sc-12762 (6E4/4) | Santa Cruz |
| anti-Six1 | Rabbit | 1:200 | #12891S (D4A8K) | Cell Signaling Technology |
| anti-Eya1 | Rabbit | 1:100 | #PA5-65034 | Invitrogen |
| anti-Laminin | Rat | 1:200 | #ab44941 (LT-3) | Abcam |
| anti-Fibronectin | Goat | 1:300 | #sc-6953 (N-20) | Santa Cruz |
| anti-Snai1 | Rabbit | 1:100 | #C15D3 | Cell Signaling Technology |
| anti-mCherry | Rat | 1:400 | #M11217 | ThermoFisher |

| | | | | |
|---|---|---|---|---|
| Phalloidin AF488 | | 1:1000 | #A12379 | ThermoFisher |
| Phalloidin AF635 | | 1:1000 | #A34054 | ThermoFisher |

**Table 2: List of primers used for RT-PCR**

| Gene | Forward Primer | Reverse Primer |
|---|---|---|
| Shh | GCGGCAGATATGAAGGGAAGA | CCAGGCCACTGGTTCATCAC |
| T/Bra | CTGGGAGCTCAGTTCTTTCG | GTCCACGAGGCTATGAGGAG |
| Gsc | AGACGAAGTACCCAGACGTG | CTGTCGTCTCCACTTGGCTC |
| FoxA2 | CATTACGCCTTCAACCACCC | GGTAGTGCATGACCTGTTCG |
| Gapdh | CGTATTGGGCGCCTGGTCAC | ATGATGACCCTTTTGGCTCC |
